# Deoxydinucleotides activate the bacterial anti-phage defense system ApeA

**DOI:** 10.64898/2026.01.26.701840

**Authors:** Jonas Juozapaitis, Arūnas Šilanskas, Audronė Rukšėnaitė, Ema Ežerskytė, Rugilė Puteikienė, Domantas Vareika, Lidija Truncaitė, Giedre Tamulaitiene, Inga Songailienė, Virginijus Siksnys, Giedrius Sasnauskas

## Abstract

Bacteria and archaea encode diverse antiviral defense systems, many of which rely on toxic effector proteins that are activated specifically upon bacteriophage infection. However, the mechanisms by which infection is recognized and coupled to effector activation remain poorly understood for most antiviral systems. Here, we focus on ApeA, a HEPN-domain antiviral protein that confers immunity through cleavage of host tRNAs within their anticodon loops. We show that ApeA proteins form large doughnut-shaped oligomers that are activated upon ligand binding in a conserved protein pocket distinct from the catalytic center. In the Ec2ApeA variant, this pocket specifically recognizes 5′-phosphorylated deoxydinucleotides that likely arise as intermediates of host genome degradation by viral nucleases, thereby enabling Ec2ApeA to achieve a broad protection profile. Together, our results reveal how small-molecule products of virus-induced host cell destruction function as signals that activate bacterial immune defenses.

## INTRODUCTION

Co-evolution between bacteria and their bacteriophages (phages) has driven the emergence of a vast array of antiviral defense systems in bacteria, as well as counteracting strategies in phages. All bacterial immunity systems must distinguish self from non-self to become activated only upon infection, since spontaneous activation would be detrimental to the cell. Activation of bacterial immunity is often triggered by specific DNA or RNA sequences (e.g., RM and CRISPR–Cas systems) or structures (e.g., cGAS-like systems) ^1^, including free DNA ends (e.g., Shedu, Kiwa) ^2,3^, or structural phage proteins (e.g., Thoeris, AVAST, CBASS) ^4–6^, and components of the phage replication machinery, like single-stranded DNA–binding proteins (SSBs) as exemplified in retrons and other systems ^5,7^. Hundreds of new antiviral defense systems have been identified recently, however, for most of them the mechanisms of activation remain unknown. In the known cases the infection either directly activates the defense protein or is amplified through the synthesis of small nucleotide-based signaling molecules, such as cyclic oligoadenylates, cyclic di/trinucleotides, 3′,5′-cyclic mononucleotides, and NAD⁺ derivatives^8^, that are subsequently recognized by the effector. The requirement for precise self/non-self discrimination is especially acute in abortive infection (Abi) systems, where a toxin such as a DNase, RNase, NADase, or membrane pore-forming protein inhibits cellular metabolism or induces programmed cell death, thereby preventing phage propagation ^9^.

Here, we focus on ApeA, one of the first identified antiphage systems that contain a HEPN (<u>h</u>igher <u>e</u>ukaryotes and <u>p</u>rokaryotes <u>n</u>ucleotide-binding) RNase domain ^10,11^. HEPN proteins are widespread across all domains of life and typically form dimers with a composite active site at the subunit interface ^12^. In bacterial defense, HEPN acts as the RNase effector domain of Cas13 ^13^, CRISPR-accessory proteins Csm6 ^14^ and Csx1 ^15^, anticodon nuclease PrrC ^16^, and other systems ^11,17–19^. Unlike most multicomponent systems, ApeA consists of a single protein harboring a C-terminal HEPN domain and an N-terminal region with no detectable homology to known proteins ^11,12^. Here we aimed to establish the mechanism of antiviral defense provided by ApeA using a combination of in vivo, biochemical, and structural analyses.

We show that ApeA proteins form large, symmetric, doughnut-shaped oligomeric assemblies that, upon activation, cleave tRNA at the anticodon loop. We further demonstrate that ApeA’s RNase activity is triggered through ligand binding within a conserved pocket formed by the HEPN and N-terminal domains. In the case of the Ec2ApeA variant, activation occurs upon binding of 5′-phosphorylated deoxydinucleotide fragments with a 3′-terminal guanosine (pdNG), which are likely generated during host genome degradation by viral nucleases. This enables Ec2ApeA to achieve a broad protection profile, and establishes small-molecule products generated during virus-induced host cell destruction as a novel class of activity triggers in bacterial immunity.

## RESULTS

### ApeA is an abortive infection system

Using the first characterised ApeA protein sequence ^11^ as a query, we performed PSI-BLAST searches to identify homologous proteins and constructed a maximum-likelihood phylogenetic tree. From the resulting set, we selected five additional ApeA homologs sharing 20–60% sequence identity with the original sequence, prioritizing systems from *E. coli* or related species to match our heterologous host *E. coli*, but also including a homolog from the thermophilic actinomycete *Thermoactinospora rubra* (Figure 1A, Supplementary file S1,2). We named the original, experimentally characterised protein ^11^ as Ec1ApeA and five additional homologs as Ec2ApeA, Ec3ApeA, Ec4ApeA, Pa1ApeA and Thr1ApeA (Figure 1A-B) respectively. Synthetic DNA fragments encoding these proteins were cloned into a low-copy pACYC vector under an IPTG-inducible T7 promoter.

**Figure 1.**
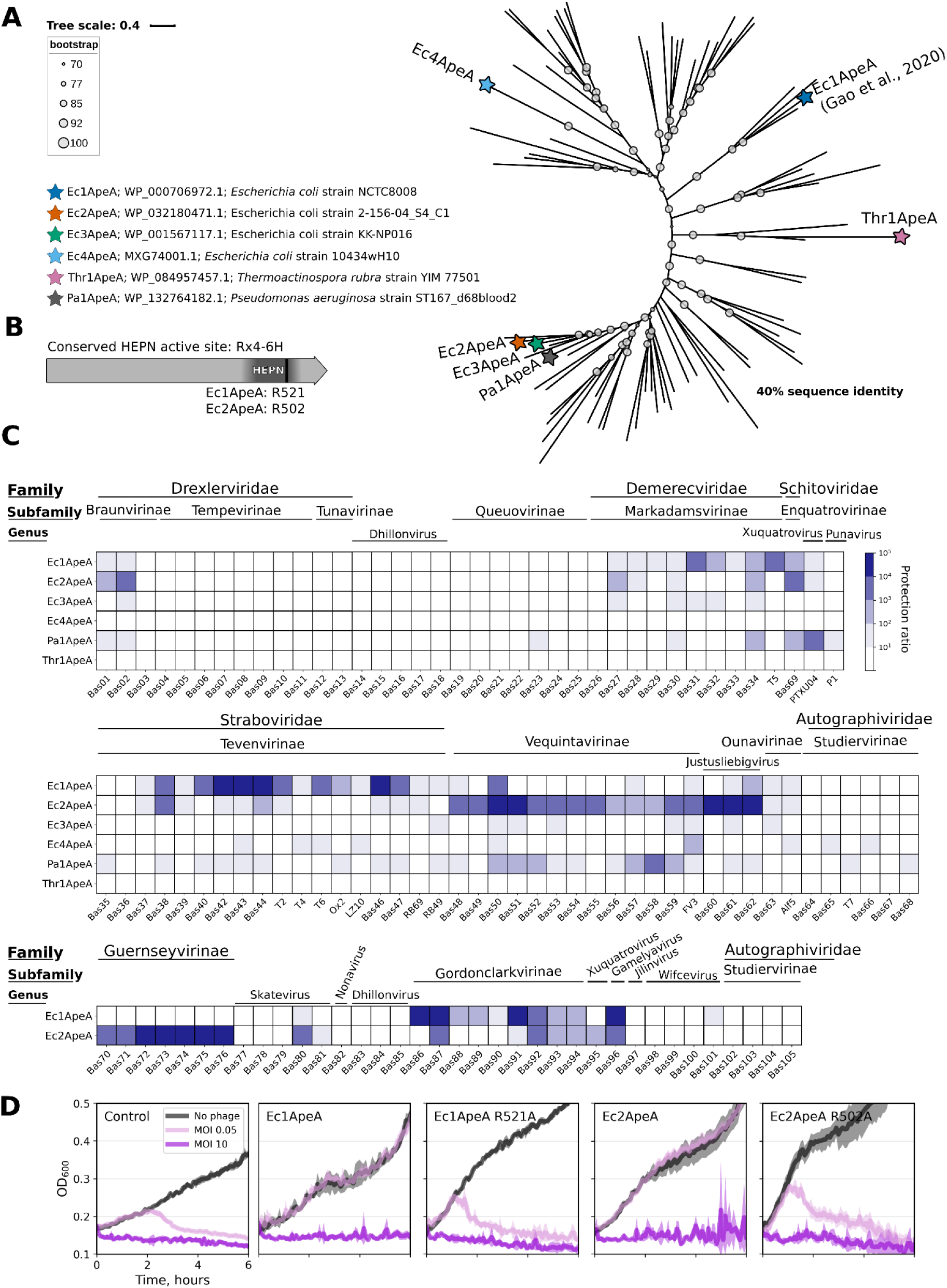
Selection and antiviral activity of ApeA proteins. (A) Maximum likelihood phylogenetic tree of ApeA protein sequences clustered at 40% sequence identity. Proteins selected for experimental analysis are marked by stars. NCBI protein IDs and bacterial strains of the selected homologs are indicated. (B) Schematic representation of ApeA defense system consisting of a single protein gene. The predicted HEPN domain is shaded in dark grey. (C) Plaque assay of *E. coli* expressing different ApeA proteins challenged with a set of *E. coli* bacteriophages. Heatmap represents the log10 of the protection ratio. (D) Growth of *E. coli* BL21 (DE3) carrying the empty vector or the plasmid-encoded Ec1 or Ec2 ApeA variant during the infection of phage RB69 at low and high MOI. Ec2ApeA R502A and Ec1ApeA R521A HEPN catalytic center mutations inactivate the defense. The shaded area depicts the standard deviation from three replicates.

To assess the antiviral activity of the ApeA homologs, *E. coli* strains carrying the corresponding plasmids were challenged with a broad panel of bacteriophages, including classical T-even phages, phages from the BASEL ^20,21^, and in-house *E. coli* phage collections (Supplementary File S1). Among tested variants, Ec1ApeA and Ec2ApeA conferred the strongest protection, Ec3ApeA, Ec4ApeA, and Pa1ApeA provided moderate protection, whereas Thr1ApeA showed no detectable antiviral activity in *E. coli* (Figure 1C). As expected, activity of Ec1ApeA and Ec2ApeA proteins was abrogated by HEPN active site R521A and R502A mutations respectively (Figure 1B,D). Notably, phages of the *Tevenvirinae* subfamily were more sensitive to Ec1ApeA, whereas phages of the genera *Vequintavirus*, *Justusliebigvirus*, and *Guernseyvirinae* family were sensitive to Ec2ApeA. The remaining active systems did not exhibit any pronounced specificity toward a particular taxonomic group under the tested conditions (Figure 1C). In contrast, phages of the *Autographiviridae* family, the *Queuovirinae* subfamily, and the *Dhillonvirus* genus were resistant to ApeA-mediated defense. These data indicate that ApeA provides broad protection against specific phage groups and that the spectrum of sensitivity varies among ApeA homologs (Figure 1C). During the plaque assays we were unable to detect any escaper phages for either of the ApeA systems.

Next, we challenged Ec1ApeA and Ec2ApeA systems with phage RB69 in liquid culture at both low and high multiplicities of infection (MOI = 0.05 and 5). Phage RB69, which is characterized by a rapid lysis phenotype ^22^, was chosen to facilitate observation of bacterial culture collapse. Both ApeA homologs confer protection at low MOI but not at high MOI, when most cells are infected (Figure 1D). These results indicate that ApeA functions via an abortive infection mechanism ^17^.

### ApeA confers antiviral protection via tRNA anticodon cleavage

To identify RNA targets of ApeA, we infected exponentially growing *E. coli* cells expressing Ec1ApeA or its catalytic center mutant with the well-characterized phage T2 at a high MOI, and extracted total RNA at different time points during infection (Figure 2A). Urea-PAGE analysis revealed extensive RNA degradation in cells expressing wild-type Ec1ApeA, but not the mutant variant (Figure 2B). Since RNA degradation products were most prominent 60 minutes after infection, we performed RNA sequencing on samples collected at that time point from both wild-type and mutant-expressing cells. To precisely map RNA cleavage sites, we used an adapter ligation–based RNA-seq library preparation method, allowing us to detect the positions of newly formed RNA ends specific to the active RNase sample (Figure 2C, Supplementary File S1), as previously described ^23^. The majority of genomic positions, where newly formed RNA ends were enriched at least 100-fold relative to the control (Supplementary Table S2), mapped to tRNA genes, with most cleavages occurring within the anticodon loop (Figure 2C). The strongest signal was observed for Pro, Thr, Met, and Ser tRNAs (Figure 2D, Supplementary File S1). As expected, no enrichment of tRNA cleavage products was observed in the sample obtained with the HEPN mutant (Supplementary File S1). Together, these results demonstrate that Ec1ApeA exhibits endoribonuclease activity targeting the tRNA anticodon loop, which is specifically activated during bacteriophage infection.

**Figure 2.**
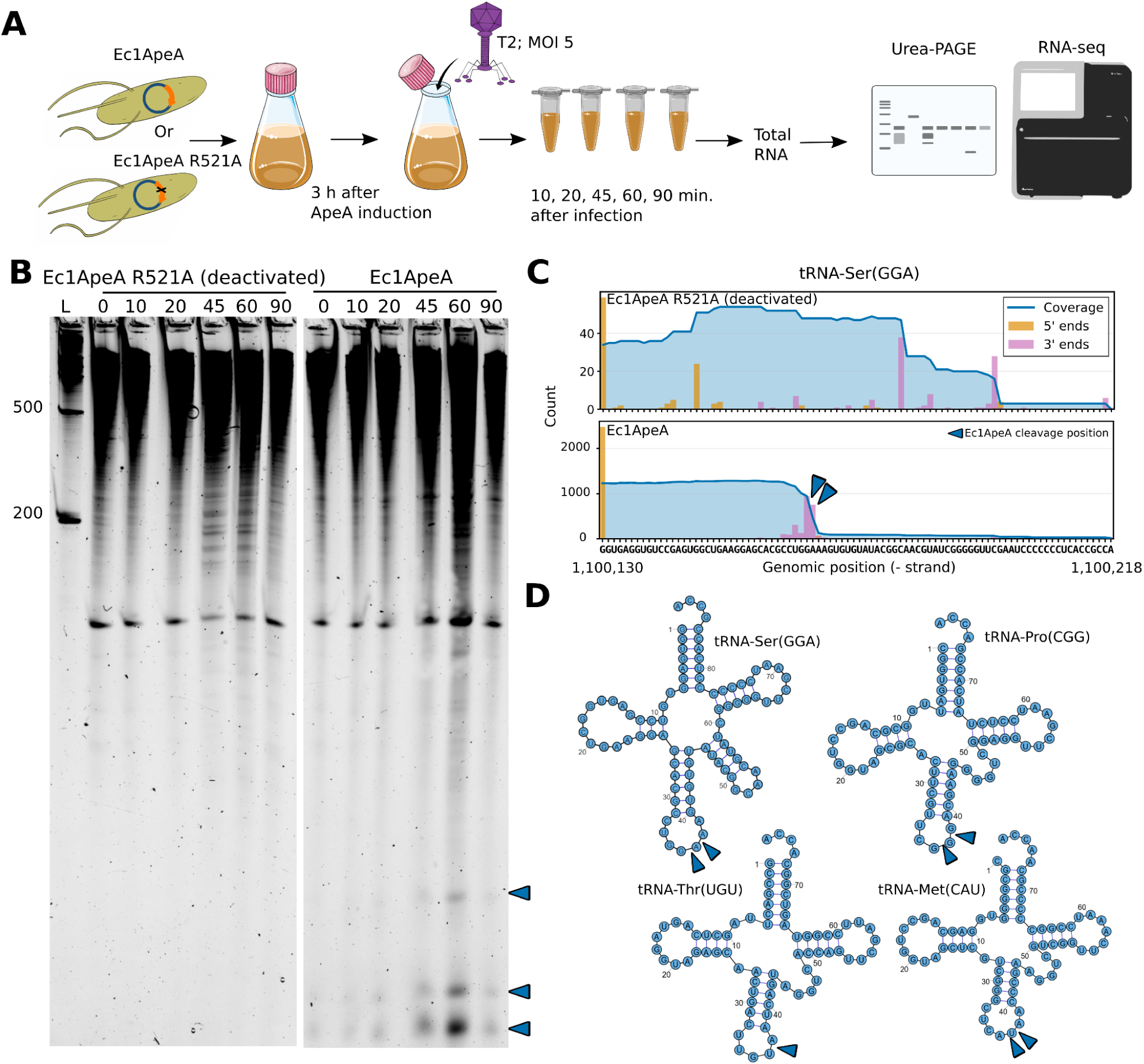
Identification of the ApeA ribonuclease target. (A) Experimental workflow for Ec1ApeA RNA preparation and cleavage target identification. (B) Urea-PAGE analysis of total RNA of *E. coli* expressing wild-type and deactivated Ec1ApeA proteins. (C) Example of tRNA cleavage position in tRNA-Ser(GGA) identified by tracking RNA-seq read end enrichment in the active-RNase case. (D) Representation of predicted Ec1ApeA cleavage positions detected by the highest read-end enrichment in the analysis. Cleavage positions are marked by blue triangles. 2D RNA sketches represent the general secondary structures of bacterial tRNAs ^40^.

### Overall structure of ApeA proteins

Ec2ApeA, Ec4ApeA, and Thr1ApeA proteins carrying C-terminal His-tags were purified by Ni^2+^ affinity chromatography followed by size-exclusion chromatography. The less soluble Ec1ApeA and Ec3ApeA proteins were purified as fusions with an N-terminal MBP-Twin-strep-10His-TEV “supertag” ^24,25^. Tagging the proteins did not interfere with the anti-phage in the cell (Supplementary Figure S1A). As revealed by SEC-MALS analysis, ApeA proteins form dimeric and more prominent higher order oligomeric species (Supplementary Figure S1BCD).

Using Cryo-EM, we determined the structures of Ec1ApeA, Ec2ApeA, Ec3ApeA, Ec4ApeA, and Thr1ApeA. All ApeA homologs share similar monomer architecture and, as expected for HEPN-domain proteins ^12,26^, form primary dimers (Supplementary Figure S2A, Figure 3A). However, in contrast to most HEPN domain proteins characterized to date, ApeA dimers, except the Thr1ApeA, assemble into larger, symmetric doughnut-shaped complexes: Ec1ApeA, Ec2ApeA and Ec4ApeA formed decamers (pentamers of dimers), whereas Ec3ApeA assembled into a dodecamer (hexamer of dimers) (Figure 3A). Further structural analysis is based primarily on the Ec2ApeA complex, which, in the case of the catalytic mutant R502A, was solved to the highest resolution (2.53 Å).

**Figure 3.**
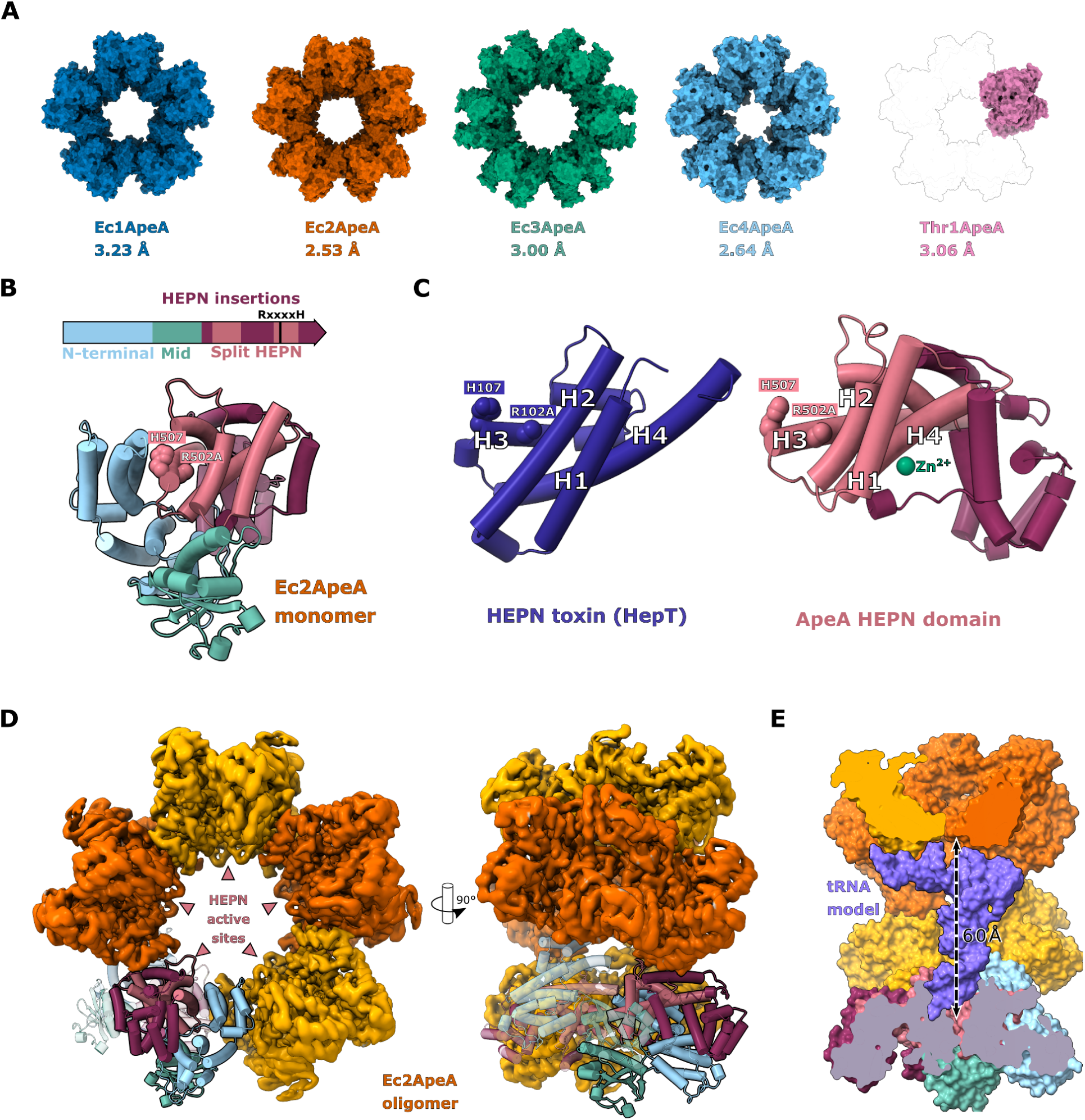
The overall structure of ApeA. (A) Comparison of ApeA protein structures determined in this study. Thr1ApeA exists as a dimer; a grey outline of a decameric structure is shown to illustrate the similarity between dimeric units. (B) *Top*: Ec2ApeA gene colored by domain boundaries. The ApeA monomer is shown with the HEPN domain in pink, the Mid domain in green, and the N-terminal domain in light blue. HEPN domain insertions absent in canonical HEPN structures ^12^ are colored purple. *Bottom*: Structure of the ApeA monomer, with HEPN active site residues marked by pink spheres. (C) Comparison of the HEPN (HepT) toxin (PDB ID: 7AE8) and the Ec2ApeA HEPN domains. Canonical HEPN ɑ-helices (H1-H4) are numbered as described in ^12^. Conserved HEPN active site residues are shown as spheres. In both structures, the active site arginine is mutated to alanine. The Zn^2+^ ion is depicted as a green sphere. (D) Cryo-EM map of Ec2ApeA decamer shown in two orientations. The density is colored in alternating yellow and orange to indicate individual ApeA dimers. The atomic model of one of the dimers is shown in cartoon representation, with one subunit opaque and the other transparent. (E) Surface representation of Ec2ApeA, cut through the center of the oligomer, with a model of bacterial tRNA^Gln^(CUG) manually fitted into the central channel.

ApeA monomers adopt a predominantly ɑ-helical, tightly folded structure that can be divided into three domains: N-terminal, Mid, and HEPN_ApeA (Figure 3B, Supplementary Figure S2A). The HEPN_ApeA part (Ec2ApeA region 335-583) represents a structurally diverged split HEPN domain that retains four core ɑ-helices (H1-H4) (Figure 3C). The closest structural homolog identified by GT-align ^27^ is the HepT toxin (PDB IDs:5YEP and 7AE8), which represents the minimal HEPN domain ^23,28^. The HEPN_ApeA domain contains additional insertions before H1, between H3 and H4, and after H4 ɑ-helices (Figure 3BC). The active site residues are positioned within the conserved region of helix H3 (Figure 3C), similar to other HEPN domains ^12^. The HEPN active sites in the doughnut-shaped assemblies are facing the inner channel (Figure 3D), which is roughly similar in size to the target tRNA molecule (Figure 3E). The N-terminal domain of Ec2ApeA (residues 1-198) shows structural similarity to Catenin alpha-1 protein (PDB ID: 6O3E) ^29^, whereas the Mid domain (residues 199-309) resembles the *Corynebacterium glutamicum* ubiquitin-like protein ligase PafA (PDB ID 4BJR) (Figure S2B) ^30^. Despite distant similarities, both domains likely perform distinct functions, as they correspond only to fragments of the above proteins, pack tightly against each other and the HEPN_ApeA domain, whereas the Mid domain lacks the catalytic residues characteristic of PafA.

ApeA monomers dimerize through the conserved core of the HEPN_ApeA domains, forming a composite HEPN active site, and through additional contacts between Mid and HEPN_ApeA domains (Figure 4A). Ec2ApeA and Ec3ApeA also coordinate Zn^2+^ ions at the primary dimerization interface between the HEPN_ApeA domain of one subunit and the Mid domain of the partner subunit (Figure 3C, Supplementary Figure 2C). These Zn^2+^ sites are not conserved in other homologs, suggesting a primarily structural role.

**Figure 4.**
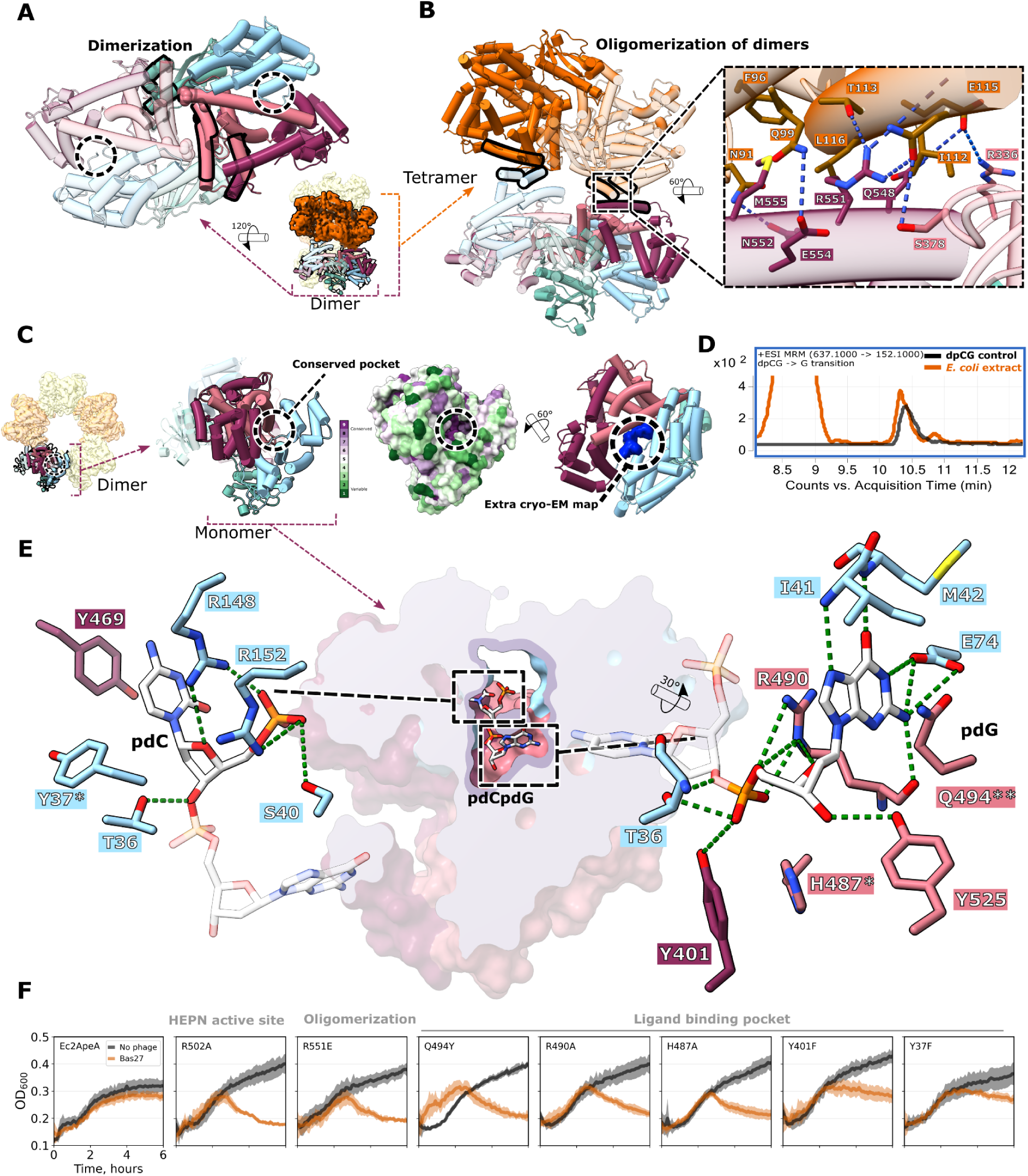
Key features of Ec2ApeA structure. (A) Structure of Ec2ApeA primary dimer, with one subunit opaque and the other transparent. Domains are colored as in Figure 3. HEPN active-site residues are depicted as pink spheres. The ɑ-helices of one of the subunits involved in dimerization are highlighted in black, and the conserved pocket formed by the HEPN and N-terminal domains is marked by dashed circles. (B) *Left*: Zoom-in view of two adjacent Ec2ApeA primary dimers. The ɑ-helices of both subunits involved in oligomerization of primary dimers are highlighted in black. *Right*: Zoomed-in view of the oligomerization interface, with hydrogen bonds shown as blue dashed lines. (C) Unmodeled density in the conserved Ec2ApeA pocket (marked by a dashed circle). *Right*: Extra density not connected to the protein is shown in blue. (D) Mass spectrometry analysis of *E. coli* extracts showing identification of the dpCG dinucleotide in exponentially growing cells. Synthetic dpCG nucleotide was used as a control. (E) Surface representation of the Ec2ApeA monomer saturated with dpCG, clipped through the ligand-binding pocket. Hydrogen bonds are marked by green dashed lines. Conserved amino acids not forming hydrogen bonds but confirmed as essential by mutagenesis are marked with *. Q494 is marked by **; its substitution with a bulkier tyrosine abolished ligand binding. (F) Growth curves of WT and mutant Ec2ApeA variants during infection with phage Bas27 at MOI 0.05. Shaded areas depict the standard deviation from four replicates.

Higher-order oligomerization of the primary dimers into doughnut-shaped complexes occurs via the secondary, smaller, and less conserved interfaces formed by the sides of the HEPN insertions and the N-terminal domains (Figure 4B). In Ec2ApeA, inter-dimer contacts involve hydrogen bonding among residues R551, E554, Q99, Q548, R336, and E115 in a hydrophobic patch formed by F96, L116, and I112 (Figure 4B).

### The ligand-binding pocket

All ApeA homologs tested contain a conserved surface pocket distal from the HEPN active site and formed by the N-terminal and HEPN_ApeA domains (Figure 4A,C). In the cryo-EM map of Ec2ApeA, this pocket contained additional unmodeled density that was not connected to the protein (Figure 4C). Mass spectrometry of the denatured Ec2ApeA sample revealed a 637.11547 m/z signal (Supplementary Figure S2D), consistent with a phosphorylated cytidine-guanosine deoxyribodinucleotide (dpCG or dpGC). The density is consistent with dpCG, as the hydrogen bonding pattern favors the guanine base at the 3′ end of the dinucleotide facing the bottom of the pocked and the 5′ phosphate group pointing toward the entrance. No comparable density was detected in the corresponding pockets of other ApeA homologs.

These results suggest that Ec2ApeA protein specifically binds phosphorylated dCG deoxydinucleotides. We hypothesized that a small amount of such dinucleotide is naturally present in *E. coli* cells during normal growth, presumably formed as byproducts of DNA damage and repair pathways ^31^, and then co-purifies with Ec2ApeA. Indeed, dpCG was detectable in mass spectra of cell extracts from exponentially growing *E. coli* cultures (Figure 4D). Addition of synthetic pdCG to purified Ec2ApeA increased its melting temperature by approx. three degrees, as measured by differential scanning fluorimetry (DSF), further supporting specific ligand binding (Supplementary Figure S2E).

A cryo-EM structure of Ec2ApeA saturated with dpCG confirmed specific binding of the dinucleotide within the pocket. The 3′-terminal guanine base forms base-specific H-bonds with E74 and the main chain atoms of M42 and I41, whereas the 5′-terminal cytosine base forms no base-specific contacts, but is sandwiched between the side chains of R148 and Y469 (Figure 4E). The interaction is further stabilized by residues R148, R152, R490 T36, S40, Y401, and Y525, which form salt bridges, H-bonds and van der Waals contacts with the phosphate and deoxyribose moieties (Figure 4E).

### Oligomerization and ligand binding are critical for antiphage defense

To assess the importance of oligomerization and ligand binding in ApeA antiviral activity, we generated a set of mutants based on the structure of ligand-bound Ec2ApeA. Disruption of the oligomerization interface by the charge-reversal mutation R551E abolished Ec2ApeA decamer formation in solution (Supplementary Figure S1CD). The cryo-EM structure of the R551E mutant, resolved to 3.56 Å resolution, revealed an intact dimeric assembly with minor positional shifts between the monomers (Supplementary Fig S2F). The R551E mutant no longer protected bacteria from phage infection (Figure 4F), but still co-purified with bound dpCG, as confirmed by mass spectrometry (Supplementary Figure S3A), and was also stabilized by the ligand in the DSF assay (Supplementary Figure S3B).

Mutations within the Ec2ApeA ligand-binding pocket, Q494Y and R490A, also abolished anti-phage activity *in vivo* (Figure 4F). The Q494Y and R490A mutants no longer co-purified with the dpCG dinucleotide (Supplementary Figure S3A) nor were stabilized by supplying the ligand *in vitro* (Supplementary Figure S3B). Additional pocket mutations H487A, Y401F, and Y37F, which target conserved residues not directly involved in hydrogen bonding to dpCG, similarly impaired antiviral activity and ligand binding (Figure 4F, Supplementary Figure S3B). Nevertheless, Ec2ApeA pocket mutants retained the decameric architecture (Supplementary Figure S1CD).

Together, these results indicate that both higher-order oligomerization and an intact ligand binding pocket are essential for Ec2ApeA antiviral function. We further observed that overexpressed pocket and oligomerization mutants exhibited markedly lower toxicity to *E. coli* cells compared to WT Ec2ApeA, approaching the low toxicity of HEPN catalytic site mutant R502A (Figure 4F). The reduced toxicity of pocket mutants suggests that WT Ec2ApeA can be spontaneously activated in *E. coli* through ligand binding even in the absence of phage infection, consistent with the presence of a small amount of endogenous dpCG in the *E. coli* cells (Figure 4D). Lower toxicity of the oligomerization mutant R551E further hints that ApeA is not activated via the disassembly of the decamer into dimers (Figure 4F).

### Ligand specificity of Ec2ApeA

To further characterize Ec2ApeA ligand binding specificity, we screened a panel of nucleotides using DSF. The assay revealed that the 3′-terminal guanine base is essential for binding, whereas the 5’ cytosine can be replaced by other DNA bases, with adenine yielding the greatest increase in the melting temperature (Figure 5A, Supplementary Figure S4A). Cryo-EM structures of Ec2ApeA saturated with dpAG and dpGG confirmed an almost identical binding mode, with the 5′ base forming no base-specific contacts (Supplementary Figure S4B). In DSF experiments, the 5′ phosphate was also critical for ligand binding (Figure 5A, Supplementary Figure S4C). Extending the ligand by one additional nucleotide at the 5′ end markedly reduced binding (Figure 5A, Supplementary Figure S4C). No stabilization was detected for the ribonucleotide variant pCG (Figure 5A, Supplementary Figure S4C), presumably due to the extra 2′-OH groups making exceedingly close contacts with pocket residues Y401 and H521 (Supplementary figure S4D). Collectively, these results demonstrate that Ec2ApeA is specific to pdNG deoxyribodinucleotides.

**Figure 5.**
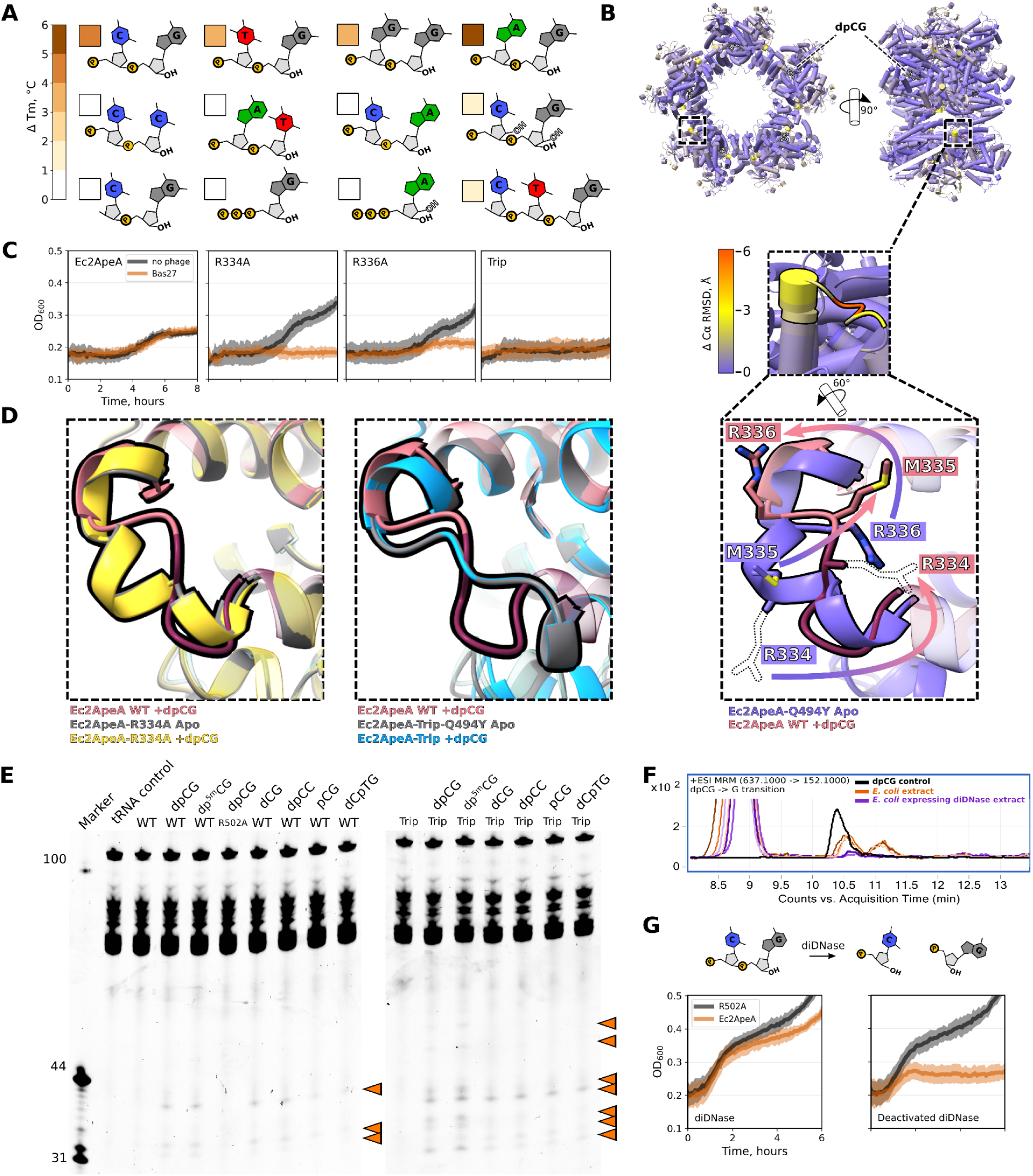
ApeA ligand specificity and activation. (A) DSF assay results shown as a heatmap, where darker shading indicates a larger change in melting temperature (ΔTm). Temperatures are compared between a control sample without ligand and the same sample with ligand added. Ligands are shown schematically, and the ΔTm upon ligand addition is indicated by the shade of the adjacent square. (B) *Top*: Conformational changes of Ec2ApeA structure induced by pdCG binding. The structure of pdCG-bound WT Ec2ApeA is colored based on C-α RMSD between the structures of pdCG-bound WT Ec2ApeA and Ec2ApeA-Q494Y proteins. *Bottom*: Zoomed-in view of conformational changes in the 331-341 aa regulatory loop of Ec2ApeA. Arrows indicate movement of R334, M335, and R336 residues from the apo-state (purple) to the ligand-bound state (pink and wine). Side chains unresolved in the cryo-EM maps are shown as dashed outlines. (C) Growth curves of Ec2ApeA regulatory loop point mutants during infection with phage Bas27 at MOI 0.05. “Trip” denotes the triple Ec2ApeA regulatory loop mutant E334A+R336E+K337A. Shaded areas represent standard deviation from 4 replicates. (D) Comparison of apo- and ligand-bound Ec2ApeA R334A and Trip mutant structures with ligand-bound WT Ec2ApeA. *Left*: Apo- and ligand-bound Ec2ApeA R334A (grey/yellow). The regulatory loop adopts the same conformation in both states, distinct from ligand-bound WT Ec2ApeA (pink and wine) but similar to inactive Ec2ApeA-Q494Y (panel B). *Right*: Apo- and ligand-bound Ec2ApeA Trip mutant (grey/blue). The regulatory loop conformation is identical in both states but differs from ligand-bound WT Ec2ApeA (brown) and Ec2ApeA-Q494Y (panel B). (E) tRNA cleavage assay of Ec2ApeA (left) and hyperactive Trip mutant (right) in the presence of various nucleotide ligands. “L” indicates the RNA ladder, Trip denotes the triple Ec2ApeA regulatory loop mutant (R334A+R336E+K337A). Orange triangles indicate Ec2ApeA cleavage products. (F) Multi-reaction monitoring mass spectrometry analysis of *E. coli* metabolites extracted from control cells and cells expressing the deoxydinucleotide-specific diDNAse enzyme ^31^. Synthetic dpCG was used as a control. Brown shades indicate 3 replicates of control *E. coli* extracts, and purple shades indicate replicates of diDNase-expressing cells. (G) Growth curves of Ec2ApeA expressing active or inactive protein variants in the background of active or inactive diDNAse. Shaded areas represent standard deviation from 4 replicates.

### Conformational changes induced by ligand binding

To gain insights into how ligand binding affects Ec2ApeA structure, we solved the cryo-EM structure of the apo Ec2ApeA Q494Y pocket mutant to 2.53 Å resolution and compared it to the dpCG-bound WT complex. The overall decameric assemblies of both variants overlay with an RMSD of less than 0.64 Å (Figure 5B, Supplementary Figure S1CD). The most notable difference involves rearrangement of the 331-341 aa loop, located near the inter-dimer interface but distal from both the HEPN catalytic center and the ligand-binding pocket. This rearrangement exposes residue R336 on the protein asurface, enabling formation of an additional inter-dimer contact (Figure 4B), while simultaneously reorienting R334 away from the surface (Figure 5B).

Protection assay in liquid medium revealed that loop mutations R334A and R336A impaired Ec2ApeA antiviral activity (Figure 5C) and reduced toxicity (Supplementary Figure S5A). Cryo-EM structures of the Ec2ApeA R334A mutant, determined in the apo (no ligand density present in the pocked) and dpCG-bound states, solved to 2.53 and 2.39 Å resolution respectively, revealed that in both cases the 331-341 loop adopts the conformation observed in the inactive pocket mutant Q494Y (Figure 5D), while the overall conformation of the complex remained unchanged (Supplementary Figure S5B). Thus, the rearrangement of the 331-341 loop upon ligand binding in the apo/ligand-bound Ec2ApeA correlates with the activity of Ec2ApeA: presumably, the R334A mutation blocks this rearrangement, thereby locking the protein in an inactive state. This interpretation is further supported by the triple loop mutant R334A+R336E+K337A (hereafter, the Trip mutant), designed to further disrupt its regulatory function. The Ec2ApeA-Trip, along with impaired antiviral activity, also displayed markedly increased toxicity compared to WT Ec2ApeA (Figure 5C, Supplementary Figure S5A). Cryo-EM structures of Ec2ApeA-Trip in the apo state (Ec2ApeA-Trip with an additional pocket mutation Q494Y to prevent ligand binding, resolution 2.63 Å) revealed a more extended 331-341 loop conformation (Supplementary Figure S5C), accompanied by translational repositioning of the adjacent ɑ-helix and a 3 Å shift of the Mid domain aa 208-304 (Supplementary Figure S5C). Ligand binding (dpCG-bound Ec2ApeA-Trip, resolution 2.86 Å) did not alter the loop conformation (Figure 5D), yet resulted in a 3 Å shift of the 210-250 aa Mid domain region located at the exterior of the decameric ring (Supplementary Figure S5D).

Taken together, these findings suggest that the Ec2ApeA 331-341 loop serves a regulatory role by mediating communication between the ligand-binding pocket and the HEPN catalytic center. However, because the conformational changes observed upon ligand binding and loop mutation do not involve the catalytic center, the molecular mechanism underlying this allosteric coupling remains to be elucidated.

### Ec2ApeA is activated by deoxydinucleotides in vitro and in vivo

*In vitro* ribonuclease assays showed that under our experimental conditions tRNA cleavage by wild-type Ec2ApeA was activated by dpCG, pd5mCpdG and some of the ligands identified as suboptimal in the DSF assay, including unphosphorylated dCG and dpCC. The ribodinucleotide pCG and trinucleotide dCTG activated Ec2ApeA to a lesser extent (Figure 5E). Activation was even more pronounced for the Ec2ApeA-Trip mutant, in which dpCG and dp5mCG triggered substantially higher activity than any other ligands (Figure 5E). RNA sequencing of the Ec2ApeA cleavage products confirmed preferential cleavage within the anticodon loops of Gln, Tyr, Val, and Gly tRNAs (Supplementary Table S3), resembling the specificity of Ec1ApeA in an infected cell. Despite reconstituted activity *in vitro*, our attempts to capture and structurally characterize the ternary complex of Ec2ApeA WT or Trip variants bound to both dpCG and tRNA have so far been unsuccessful.

Cellular levels of deoxydinucleotides are known to be markedly reduced upon overexpression of the recently characterized diDNase enzymes ^31^, which specifically bind and hydrolyze pdNN deoxydinucleotides (Figure 5F). Co-expression of WT diDNase (*Cellulosimicrobium*, WP_144679698.1), but not diDNase active site mutant (D34R+E36W), substantially alleviated the background toxicity caused by Ec2ApeA overexpression (Figure 5G), confirming that endogenous deoxydinucleotides are indeed responsible for the background activation (and the associated toxicity) of Ec2ApeA in bacterial cells.

### Structural diversity of the ligand binding pockets

Residues forming the ligand-binding pocket are conserved across the ApeA family, with a sequence consensus closely resembling the Ec2ApeA pocket (Supplementary Figure S6AB). However, moderate conservation of several pocket residues (Supplementary Figure S6B) suggests that other ApeA homologs may be activated by ligands distinct from those of Ec2ApeA. A notable example is Ec1ApeA. Although the overall pocket architecture of Ec1ApeA resembles that of Ec2ApeA, it contains several key substitutions: Ec2ApeA E74, which is involved in guanine base recognition, is replaced by the shorter aspartate D75; R490 and S40, which form a side wall of the guanine-binding site, are substituted by D509 and K41, respectively; residues R148 and Y159 stabilizing the 5′ cytosine base are replaced by Y159 and K488 (Supplementary Figures 6A and S6B); and the conserved H487 is replaced by T506 (Supplementary Figure S6B). As for Ec2ApeA, we identified mutations in the Ec1ApeA ligand-binding pocket (Figure 6A), oligomerization interface, and regulatory loop that impaired antiviral activity in the cell (Figure 6E), supporting the conservation of the proposed activation/signal transduction mechanism across ApeA homologs. However, purified Ec1ApeA failed to bind any of the tested ligands in DSF assays (Figure 6B), its background toxicity was unaffected by diDNAse co-expression (Figure 6C), it was not activated *in vitro* by deoxydinucleotides (Figure 6D), and had different phage specificity (Figure 1C), indicating that Ec1ApeA responds to an activator molecule distinct from that of Ec2ApeA.

**Figure 6.**
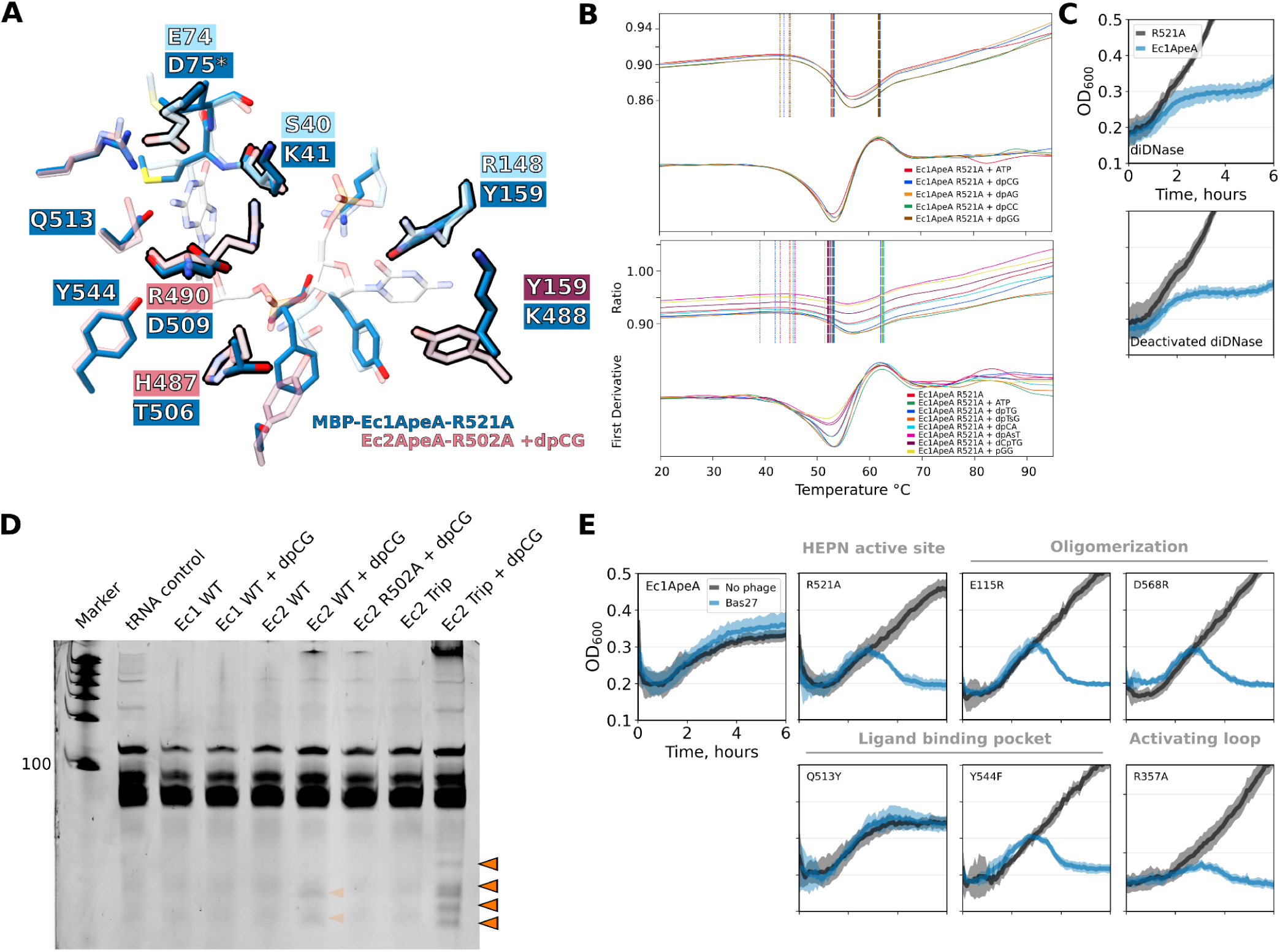
Characterization of Ec1ApeA protein. (A) Comparison of Ec2ApeA (dpCG-bound) and apo Ec1ApeA ligand binding pockets. Amino acids corresponding to Ec1ApeA are marked by dark blue boxes, and Ec2ApeA residues are indicated by pink boxes. Differences are highlighted in black. (B) DSF melting curves of MBP-Ec1ApeA incubated with different ligands. No changes in melting temperature were detected. (C) Growth curves of stains expressing active or inactive Ec1ApeA variants in the background of active or inactive diDNAse. (D) tRNA cleavage assay of Ec1ApeA and Ec2ApeA. Orange triangles indicate Ec2ApeA cleavage products. (E) Growth curves of Ec1ApeA proteins and their point mutants during infection with phage Bas27 at MOI 0.05. The shaded areas represent standard deviation from four replicates.

## DISCUSSION

In this work, we characterize the ApeA family of antiviral defense proteins. Previously, only a single functional homolog (renamed here as Ec1ApeA) had been described ^11^; here, we identify and analyze four additional active variants – Ec2ApeA, Ec3ApeA, Ec4ApeA, and Pa1ApeA. These homologs confer resistance against distinct bacteriophage taxa, revealing divergent phage specificities within the ApeA family. We further demonstrate that ApeA functions as an abortive infection system by cleaving host tRNAs within their anticodon loops to arrest translation, thereby expanding the repertoire of tRNA-targeting antiviral defense systems ^16,23,32^. Cryo-EM structures of four active ApeA homologs reveal that, in addition to primary dimers formed through HEPN domain dimerization, ApeA proteins assemble via inter-dimer interfaces into decameric or dodecameric doughnut-shaped complexes with tRNA-sized central channels. Disruption of this secondary interface in both Ec1ApeA and Ec2ApeA abolishes antiviral activity, underscoring its functional importance.

A key feature of ApeA proteins is a conserved surface pocket distinct from the catalytic center. We show that the pocket of Ec2ApeA specifically binds deoxydinucleotides (pdNG), with a preference for a 3′ guanine base, deoxyribose, and a 5′ phosphate. Mutational analyses confirm that ligand binding is essential for antiviral activity; moreover, tRNA cleavage can be reconstituted *in vitro* upon stimulation of Ec2ApeA with deoxydinucleotides. We also identify a regulatory loop (residues 331–341) in Ec2ApeA that undergoes conformational rearrangement upon ligand binding. Mutations in this loop either lock Ec2ApeA in an inactive state or cause hyperactivation, indicating that it relays the activation signal between the ligand-binding pocket and the catalytic centers. On this basis, we propose that in bacterial cells Ec2ApeA is activated upon binding of pdNG dinucleotides. Such small DNA fragments are likely generated during phage infection as intermediate products of host genome degradation by phage-encoded nucleases ^33–35^, with possible contribution from host enzymes ^36^. Notably, the phage T4 endonuclease II preferentially cleaves DNA near CG-rich sequences ^35^, potentially leading to the formation of guanosine-containing deoxydinucleotides that activate Ec2ApeA. The failure to isolate bacteriophage mutants that escape the Ec2ApeA system also supports the idea that Ec2ApeA is triggered by a general infection signal – such as host DNA degradation products – rather than by a single phage-specific component. The absence of escaper mutants further suggests that Ec2ApeA-sensitive phages may encode multiple DNases with overlapping activities involved in host genome degradation, and/or that host genome degradation is an indispensable part of their life cycle. Phages that naturally evade Ec2ApeA-mediated protection (Figure 1) may therefore differ in the extent or nature of host genome degradation during infection.

Taken together, our findings establish deoxyribodinucleotides as a novel class of antiviral defense activators, thereby expanding the repertoire of nucleotide-based molecules involved in the activation of effector proteins. Ec2ApeA also joins a broader group of HEPN-domain proteins activated by nucleotide derivatives, including Csm6 ^37^, which responds to cyclic hexaadenylate, and Csx1 ^38^, which responds to cyclic tetraadenylate. The key difference is that Ec2ApeA-activating deoxydinucleotides are generated directly during phage infection, unlike cyclic oligoadenylates that are produced by the phage RNA-sensing Csm complex ^37^. Interestingly, the recently characterized deoxydinucleotide-specific diDNAses were found to associate with antiviral defense islands ^31^, suggesting that deoxydinucleotides may play a broader role in bacterial-phage defense and antidefense strategies, with additional systems yet to be discovered.

Consistent with the distinct phage and ligand specificities of Ec1ApeA and Ec2ApeA, Ec1ApeA – and potentially other ApeA homologs – may respond to different nucleotide-based small-molecule activators that remain to be identified. Activation of ApeA proteins by small-molecule ligands may enable their artificial activation to eliminate bacterial populations or selective inhibition to enhance the efficacy of phage therapy against ApeA-producing bacteria ^39^.

## RESOURCE AVAILABILITY

### Lead contact

Requests for further information and resources should be directed to and will be fulfilled by the lead contact, Giedrius Sasnauskas.

### Materials availability

Plasmid generated in this study will be made available on request, but we may require a completed materials transfer agreement.

### Data and code availability

The EM densities and model coordinates have been deposited in the Electron Microscopy Data Bank and Protein Data Bank under the accession codes EMD-52478/PDB:9HXI (Ec2ApeA saturated with pdCpdG), EMD-52462/PDB:9HXB (Ec2ApeA Q494Y mutant), EMD-52464/PDB:9HXD (Ec2ApeA R551E mutant), EMD-52477/PDB:9HXH (Ec2ApeA R502A mutant co-purified with pdCpdG), EMD-52475/PDB:9HXG (Ec2ApeA R502A mutant saturated with pdCpdG), EMD-52468/PDB:9HXE (Ec2ApeA R502A mutant saturated with pdApdG), EMD-52474/PDB:9HXF (Ec2ApeA R502A mutant saturated with pdGpdG), EMD-56384/PDB:9TX1 (Ec2ApeA R334A mutant), EMD-56381/PDB:9TX0 (Ec2ApeA R334A mutant saturated with pdCpdG), EMD-56385/PDB:9TX2 (Ec2ApeA Trip mutant saturated with pdCpdG), EMD-56386/PDB:9TX3 (Ec2ApeA Trip and Q494Y mutant), EMD-52483/PDB:9HXN (Ec1ApeA), EMD-52482/PDB:9HXM (Ec4ApeA), EMD-52479/PDB:9HXJ (Ec3ApeA), and EMD-52481/PDB:9HXL (Thr1ApeA).

## ACKNOWLEDGMENTS

This work was supported by the Research council of Lithuania grant S-MIP-22-13 to G.S. (project leader), J.J. and I.S. were supported by the Vilnius University Fund grant MSF-JM-12/2021. I.S. was funded by the European Regional Development Fund under grant agreement number 01.2.2-CPVA-V-716-01-0001 with the Lithuanian Central Project Management Agency (CPVA).

We thank Dr. Gytis Dudas and Dr. Maria Fernanda Torres Jimenez for their consultation on phylogenetic analysis. We also thank Dr. Athanasios Typas, Dr. Jacob Bobonis, and Alessio Ling Jie Yang for screening ApeA genes using the TAC-TIC method.

## AUTHOR CONTRIBUTIONS

Conceptualization and funding acquisition – GS, JJ, IS; gene fragment design – JJ and IS. Investigation: cloning and mutagenesis – JJ and EE, Protein purification - JJ, AS, EE; SEC-MALS – JJ and AS, bacteriophage preparation – JJ, LT, EE, DV; plaque assays and growth assays – JJ, EE, DV; enzymatic assays – JJ and RP; DSF assays – JJ, Sequencing and data analysis – JJ; bioinformatic analysis – JJ, Cryo-EM – GS, RP and GT; protein structure analysis – JJ, GT and GS; mass spectrometry analysis – AR. Writing: original draft – JJ and GS, Figure preparation – JJ; consultation – VS, LT, IS, GT; review & editing – all authors.

## DECLARATION OF GENERATIVE AI AND AI-ASSISTED TECHNOLOGIES

During the preparation of this work, the author JJ used Claude.ai service in order to debug data analysis scripts. After using the service, the author reviewed and edited the content as needed and takes full responsibility for the content of the publication.

## SUPPLEMENTARY FIGURES

**Figure S1.**
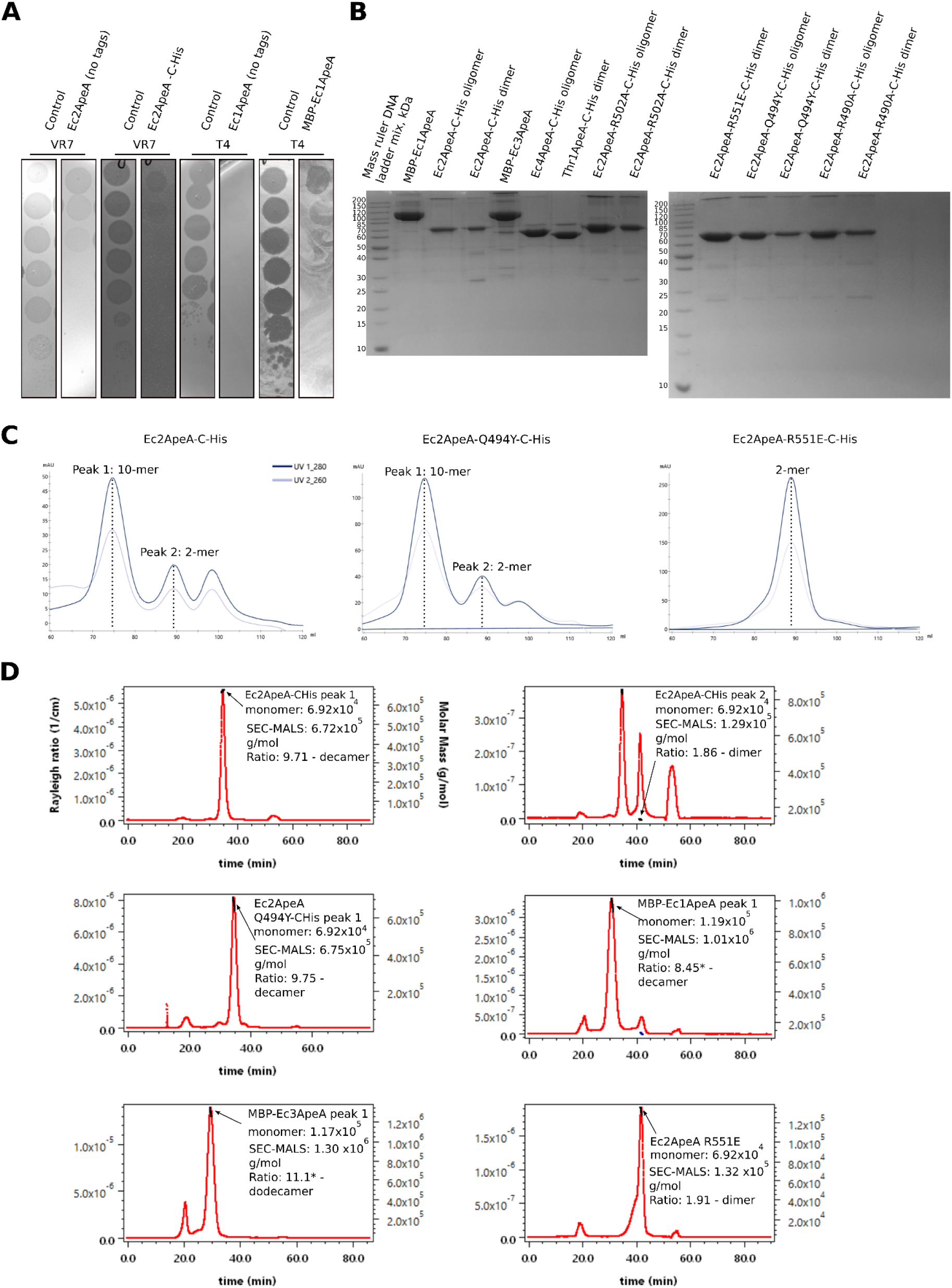
Protein purification and SEC-MALS analysis. (A) Plaque assay confirming that Ec2ApeA-CHis and MBP-Ec1ApeA tagged variants provide antiviral defense with similar efficiency to the untagged variants. Controls - empty vectors. (B) SDS-PAGE analysis of purified ApeA proteins and their mutants. (C) The final steps of ApeA purification using size exclusion chromatography. Ec2ApeA WT and Q494Y proteins eluted as two peaks (corresponding to homodecamers and homodimers), whereas the Ec2ApeA oligomerization mutant R551E eluted as a single peak (corresponding to a homodimer). (D) Size exclusion chromatography – multi-angle light scattering (SEC-MALS) analysis of different ApeA proteins. Red curves represent the light scattering signal, black lines represent the calculated particle size. “*” indicates proteins fused to the MBP-Twin-Strep-His10x-TEV supertag. The Ec2ApeA peak 2 (corresponding to a dimer, as shown in panel C), re-equilibrates after purification into a mixture of dimers and decamers.

**Figure S2.**
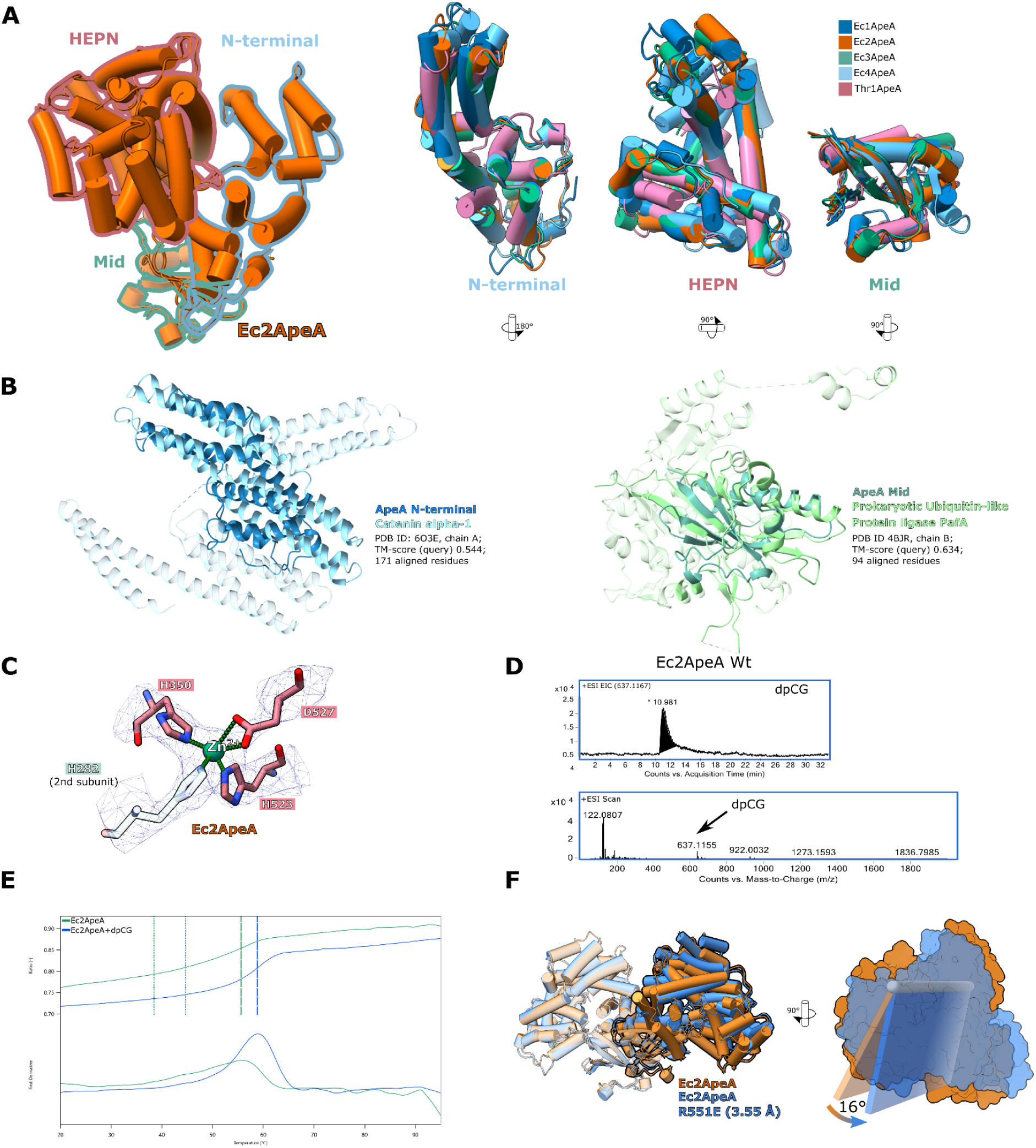
Structural details of ApeA proteins. (A) Comparison of Ec2ApeA monomer structure (*left*) with other ApeA variants. Each domain is separately aligned to Ec2ApeA (*right*). (B) Comparison of Ec2ApeA N-term and Mid domains with known protein structures identified using the GTalign search server ^41^. (C) Ec2ApeA zinc coordination site. EM density 1.9 Å around the interacting residues is depicted as blue mesh. (D) Mass spectrometry analysis of ligands bound by the WT Ec2ApeA protein. *Top*: Extracted ion chromatogram. *Bottom*: ESI scan diagram. (E) DSF curves of Ec2ApeA protein complex with and without dpCG. The ratio of tryptophan fluorescence emission at 350 and 320 nm (F_350_/F_330_) is shown along with the first derivative of this ratio, which is used for evaluation of the melting temperature. Dotted lines indicate the onset temperature (the temperature at which the protein begins to unfold). The dashed lines indicate the inflection point (the peak of the first derivative), which corresponds to the melting temperature (T_m_, the temperature at which 50% of the protein is unfolded). (F) Comparison of WT Ec2ApeA primary homodimer (part of the decameric structure) and the homodimer of the oligomerization mutant R551E (*left*). Planes depict the relative rotation of the second Ec2ApeA R551E subunit relative to the WT Ec2ApeA structure (*right*).

**Figure S3.**
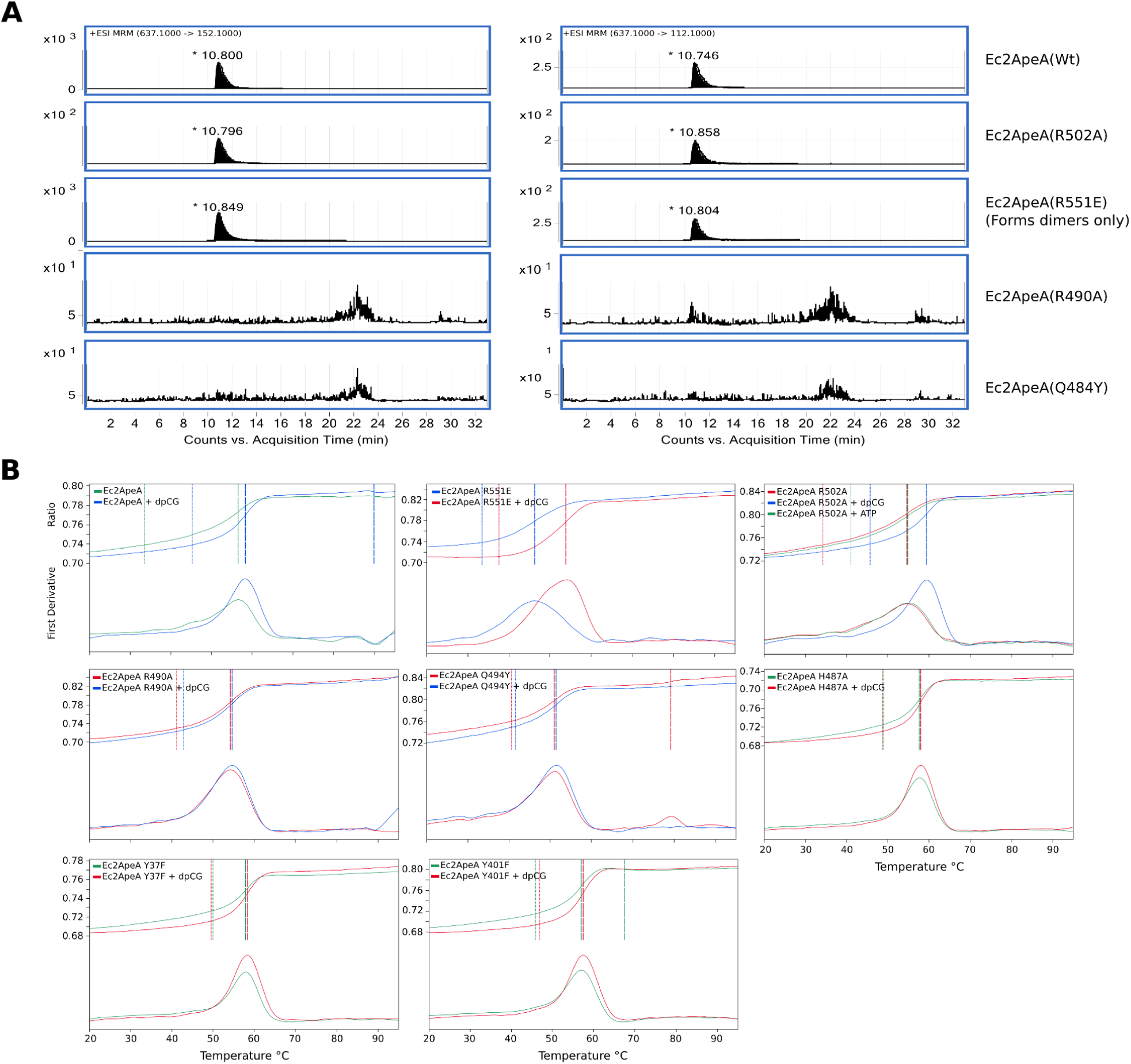
Effect of pocket and oligomerization surface mutations on ligand binding. (A) Multiple reaction monitoring (MRM) mass spectrometry analysis of ligands bound by the ApeA protein variants. The transition 637.1000 → 152.1000 corresponds to dpCG → G base transition, and 637.1 → 112.1 corresponds to dpCG → C base transition. The ligand is observed at around 10.8 min. (B) DSF melting curves of Ec2ApeA mutants with dpCG dinucleotide. ATP, used for enzymatic phosphorylation of dinucleotides, was used as a control. Ligand binding is observed for WT Ec2ApeA and the oligomerization mutant R551E, whereas binding is abolished by pocket mutations.

**Figure S4.**
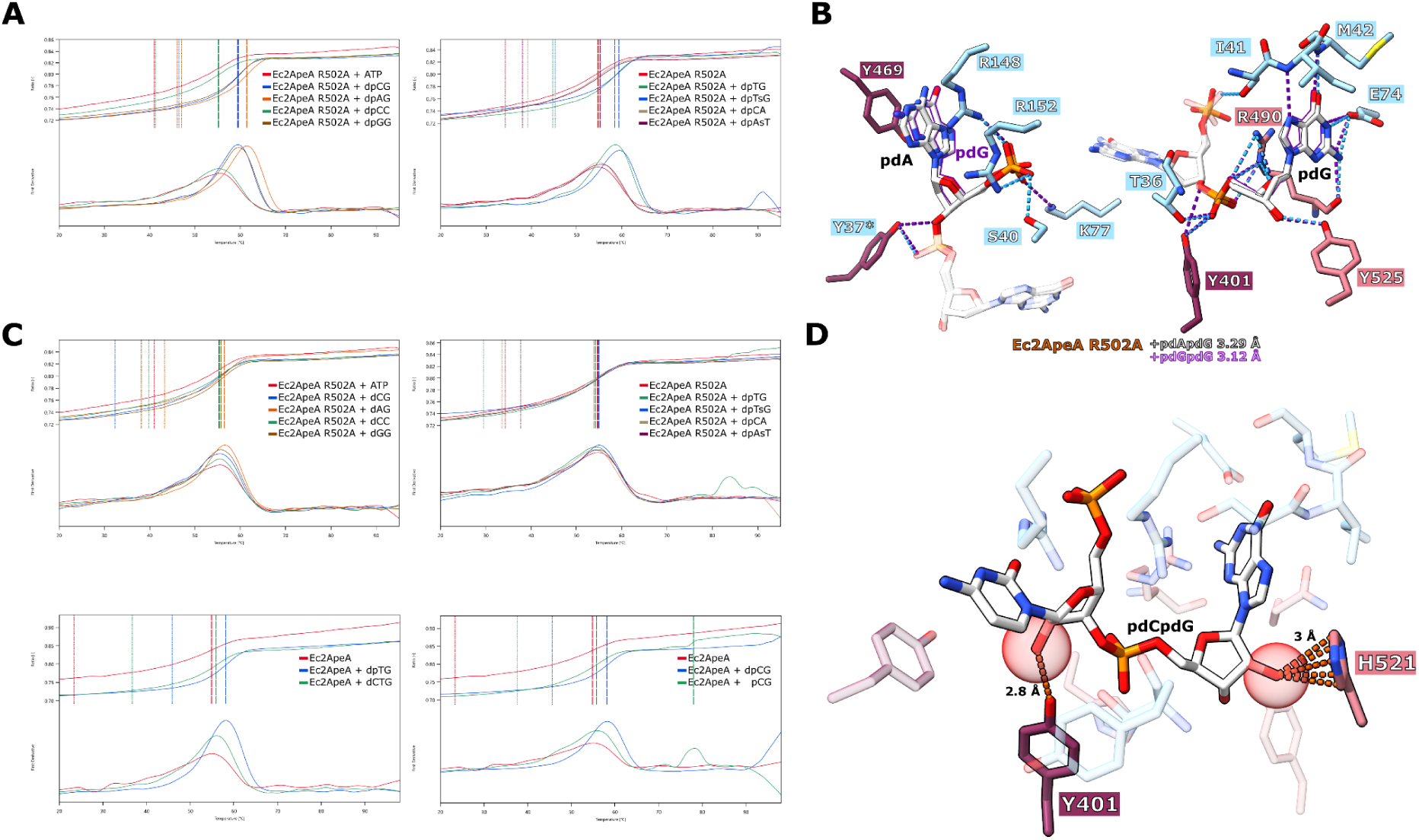
Ligand specificity of Ec2ApeA. (A) Melting curves of Ec2ApeA R502A with different 5′-phosphorylated and non-phosphorylated deoxydinucleotides. The “s” in the legend indicates a phosphorothioate modification (a non-bridging oxygen in the phosphate backbone is substituted with a sulfur atom). ATP, used for enzymatic phosphorylation of dinucleotides, was included as a control. (B) Binding of dpGG (purple) and dpAG (grey) dinucleotides in the Ec2ApeA ligand pocket. For clarity, the 5′ and 3′ nucleotides are shown separately. Contacts (<4 Å) are marked by dashed lines. (C) Melting curves of Ec2ApeA with different deoxydinucleotides lacking the 5′ phosphate, ribonucleotide pCG, and 5′-extended trinucleotide dCTG. ATP was used as a control. (D) A model of the pCG dinucleotide in the Ec2ApeA pocked based on the pdCG-bound Ec2ApeA structure. The additional 2′-OH groups are shown as red spheres. The orange dashed lines mark distances.

**Figure S5.**
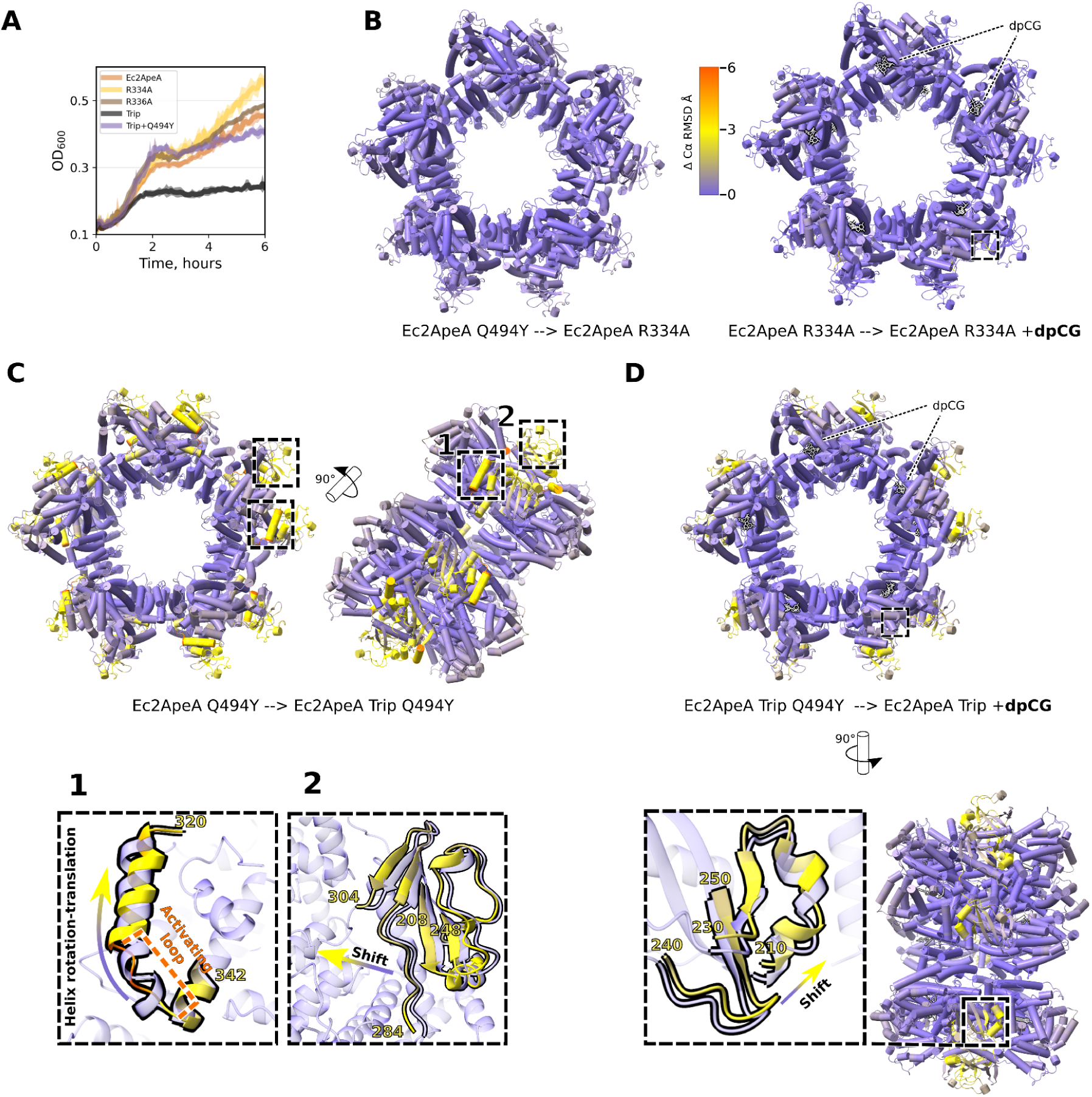
Structural effects of regulatory loop mutations in Ec2ApeA. (A) Toxicity assay of Ec2ApeA regulatory loop (residues 331-341) mutants. Growth curves of *E. coli* expressing different Ec2ApeA variants, induced with 0.1 mM IPTG, are shown. (B) *Left*: Conformational changes of Ec2ApeA structure induced by the R334A mutation. The structure of apo-Ec2ApeA-R334A is colored based on C-α RMSD between the structures of apo-Ec2ApeA-R334A and Ec2ApeA-Q494Y proteins. *Right*: changes in the Ec2ApeA R334A structure upon pdCG binding. The structure of pdCG-bound Ec2ApeA-R334A is colored based on Cɑ RMSD between the structures of apo- and pdCG-bound forms. (C) Conformational change of Ec2ApeA structure induced by the Trip (R334A+R336E+K337A) mutation. *Top*: The structure of Ec2ApeA-Trip-Q494Y is colored based on C-ɑ RMSD between the structures of Ec2ApeA-Trip-Q494Y and Ec2ApeA-Q494Y proteins. *Bottom*: Zoom-in views of the most prominent changes (region 1, the regulatory loop; region 2, residues 284-304 and 208-248). (D) Conformational change of Ec2ApeA-Trip structure induced by dpCG ligand binding. *Top*: the structure of pdCG-bound Ec2ApeA-Trip is colored based on Cɑ RMSD between the structures of pdCG-bound Ec2ApeA-Trip and apo-Ec2ApeA-Trip-Q494Y. *Bottom*: Zoom-in view of the most prominent change in the 210-230 and 240-250 aa region.

**Figure S6.**
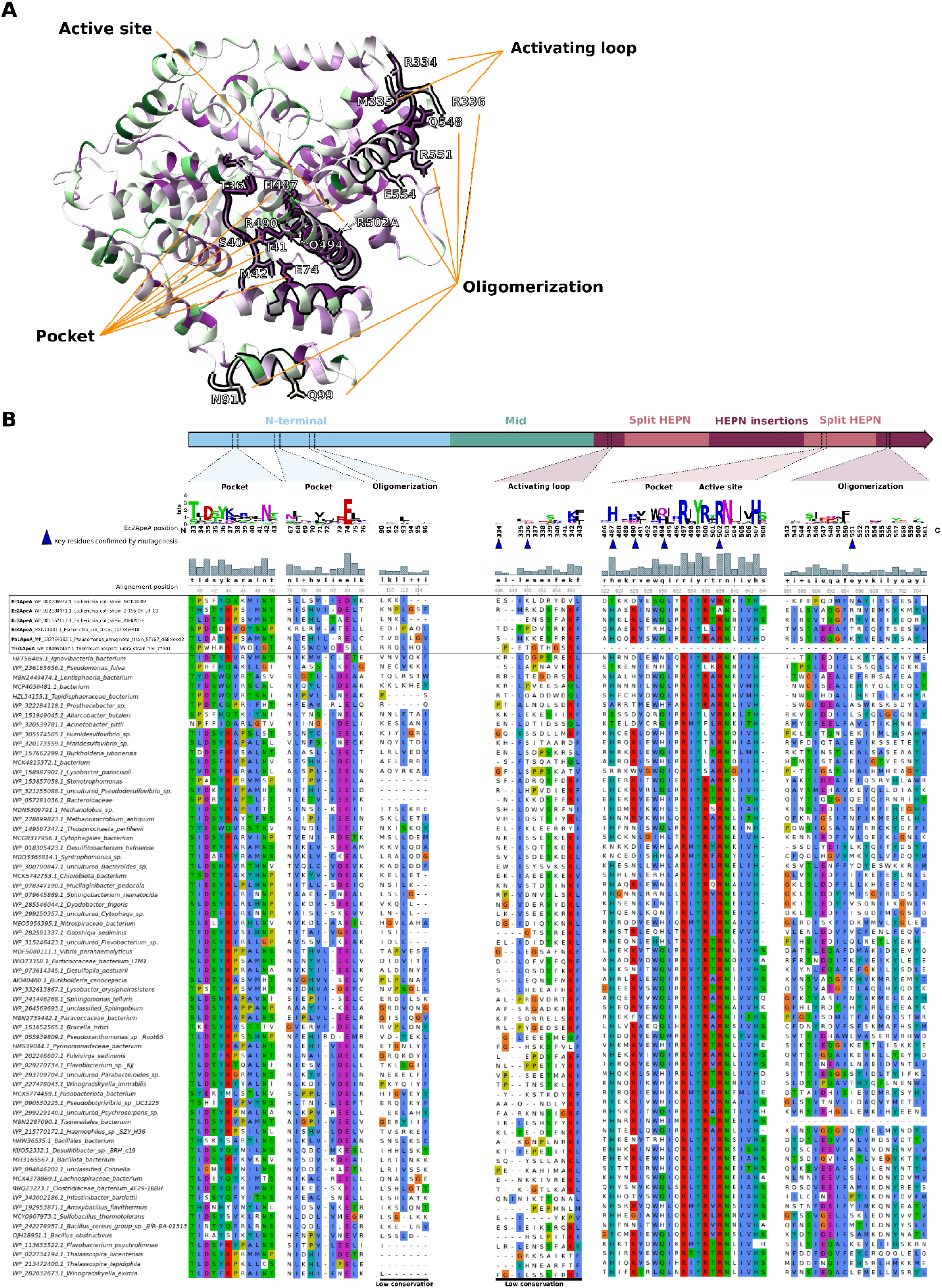
Sequence conservation of ApeA proteins. (A) Consurf conservation ^42^ analysis of the Ec2ApeA protein, using the Ec2ApeA R502A structure as a template. Functionally important protein regions are highlighted in black, with orange lines indicating their functional annotation. Key amino acids identified in this work are shown as sticks. (B) Ec2ApeA gene structure with multiple structure-based sequence alignment of functionally important protein regions. The alignment is also visualized as a WebLogo representing sequence conservation at important Ec2ApeA positions, identified in this work. Blue arrows mark key amino acids functionally confirmed by mutagenesis.

**Figure S7.**
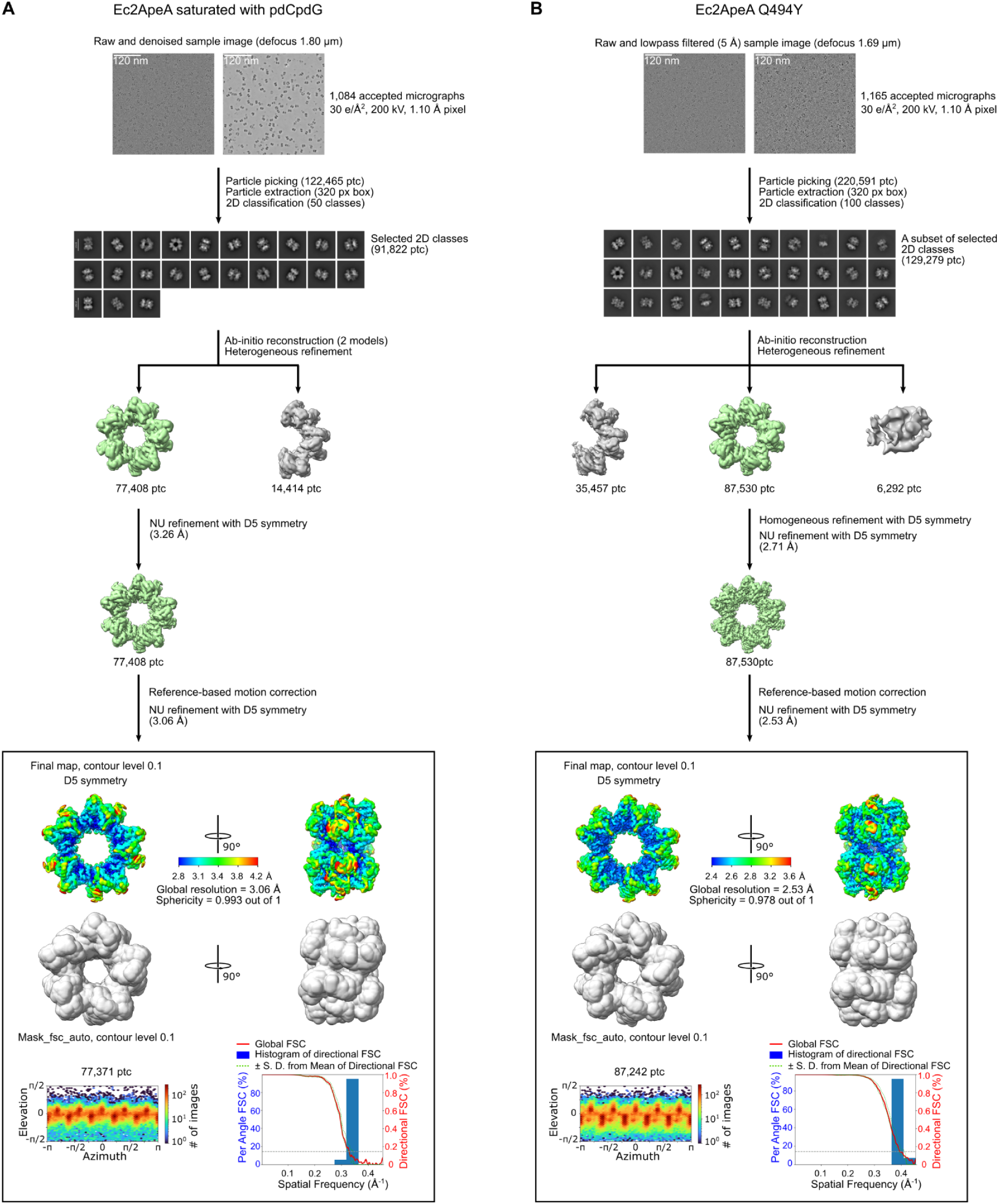
Cryo-EM data processing of Ec2ApeA saturated with pdCpdG (A) and Ec2ApeA Q494Y mutant (B).

**Figure S8.**
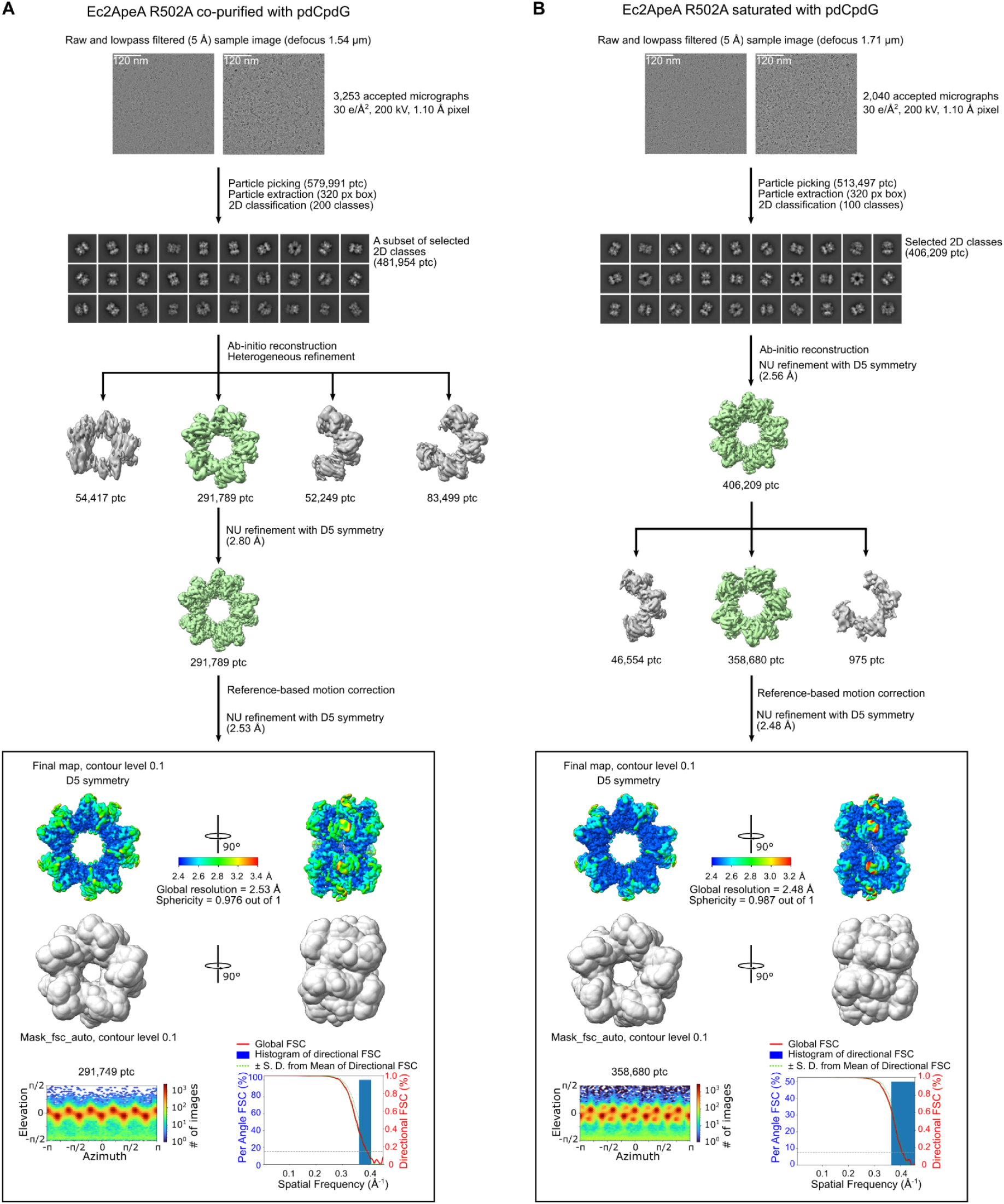
Cryo-EM data processing of Ec2ApeA R502A mutant co-purified with pdCpdG (A) and saturated with pdCpdG (B).

**Figure S9.**
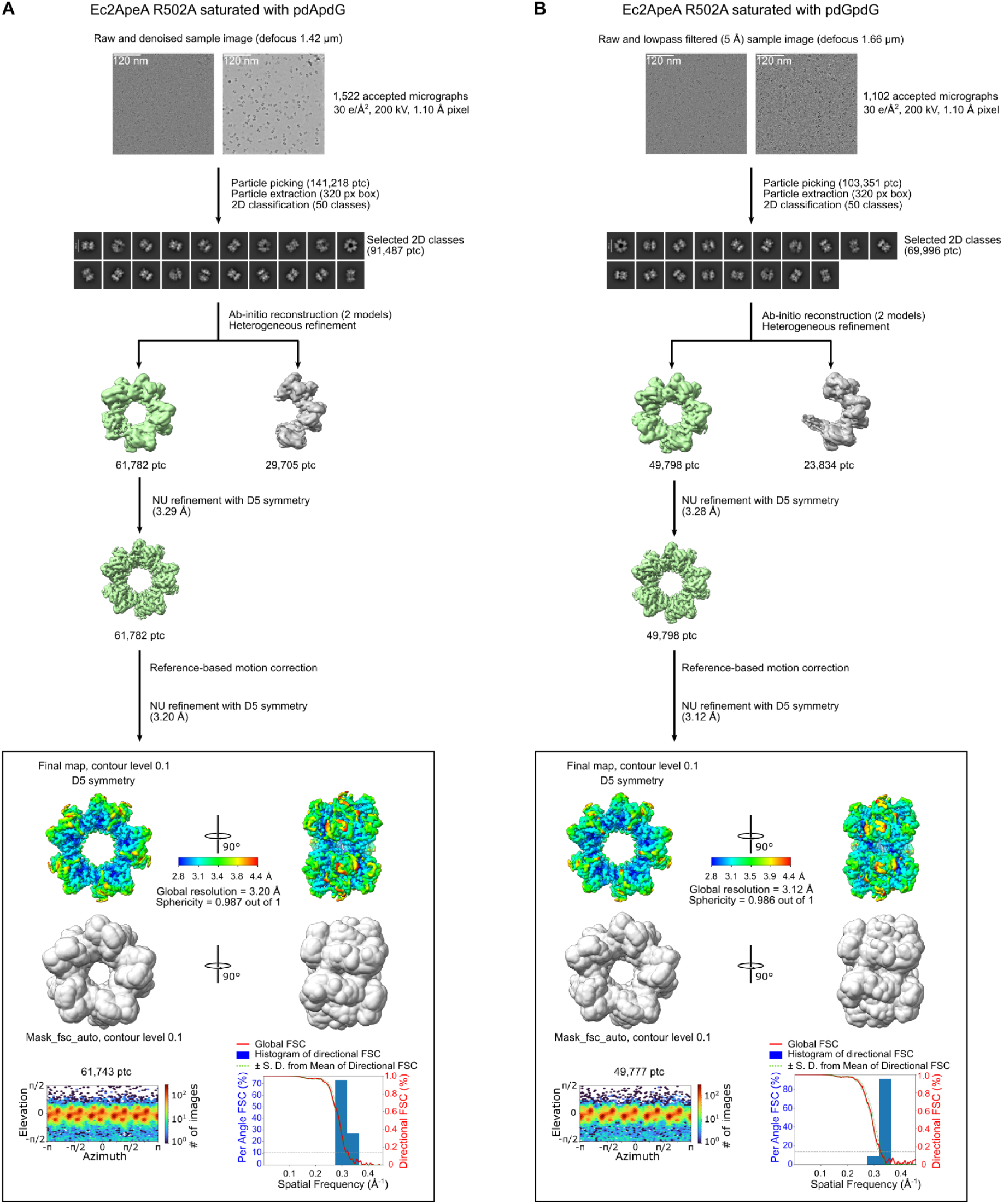
Cryo-EM data processing of Ec2ApeA R502A mutant saturated with pdApdG (A) and pdGpdG (B).

**Figure S10.**
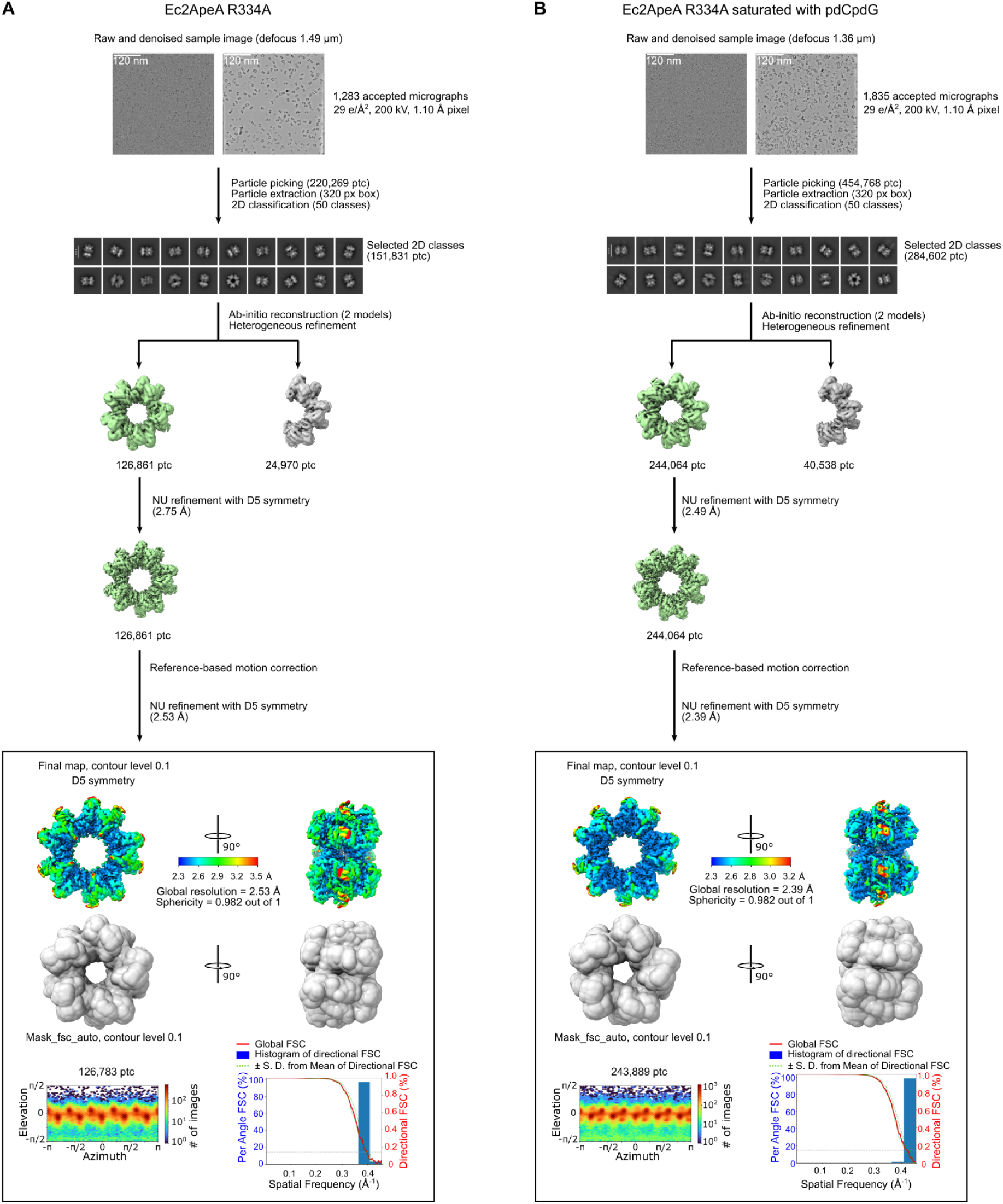
Cryo-EM data processing of Ec2ApeA R334A mutant (A) and Ec2ApeA R334A mutant saturated pdGpdG (B).

**Figure S11.**
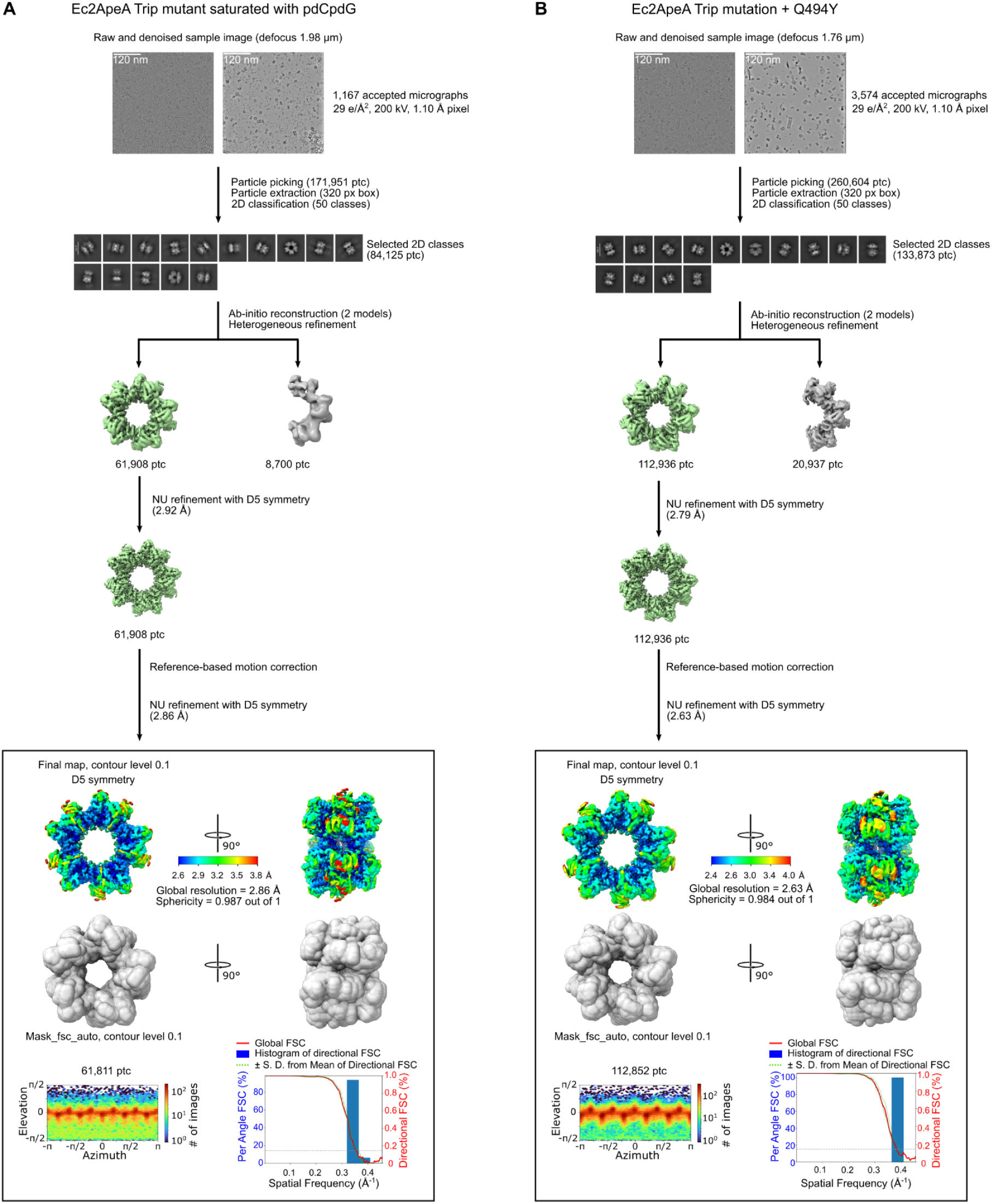
Cryo-EM data processing of Ec2ApeA Trip mutant saturated pdGpdG (A) and Ec2ApeA Trip + Q494Y mutant (B).

**Figure S12.**
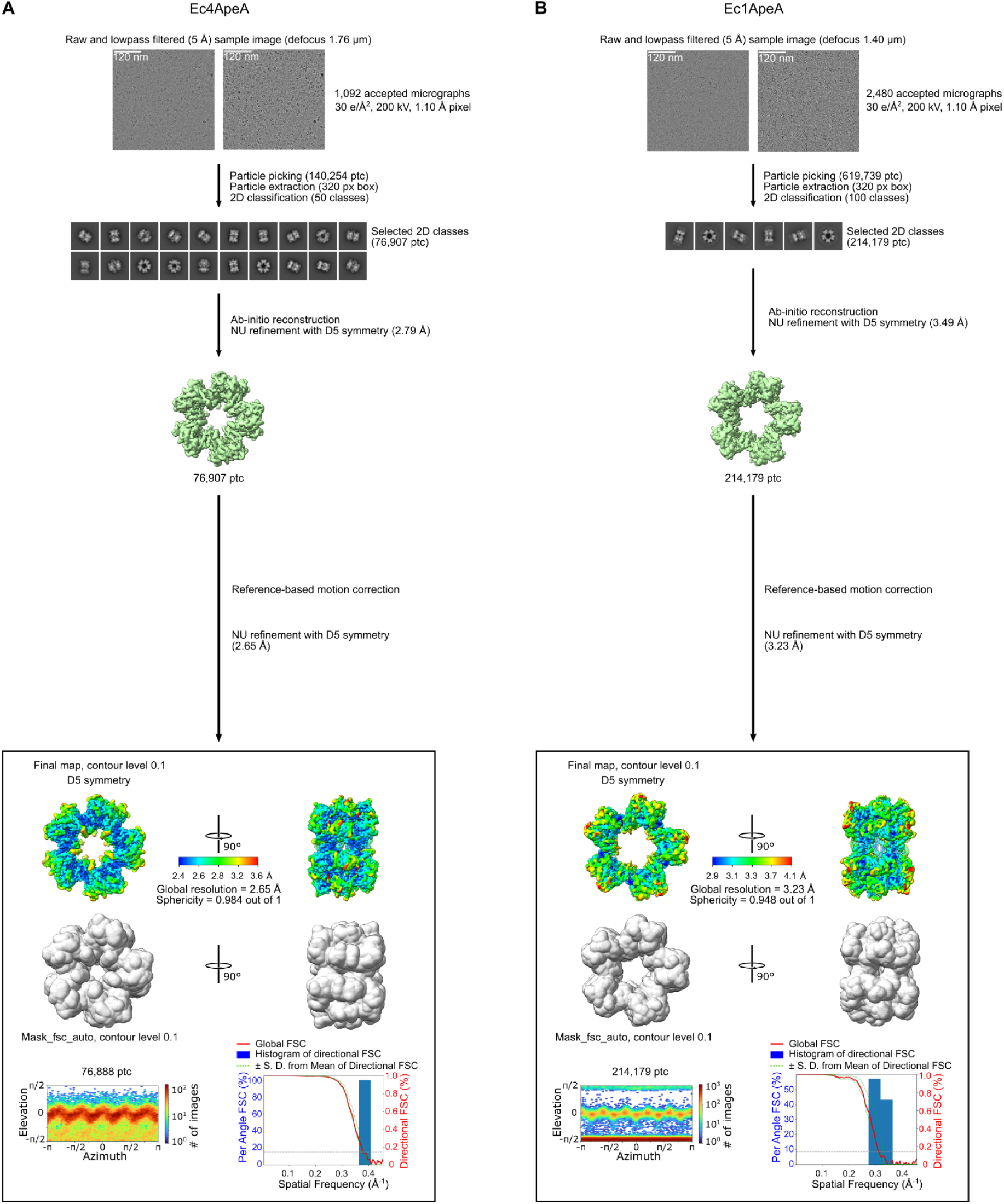
Cryo-EM data processing of Ec4ApeA (A) and Ec1ApeA (B).

**Figure S13.**
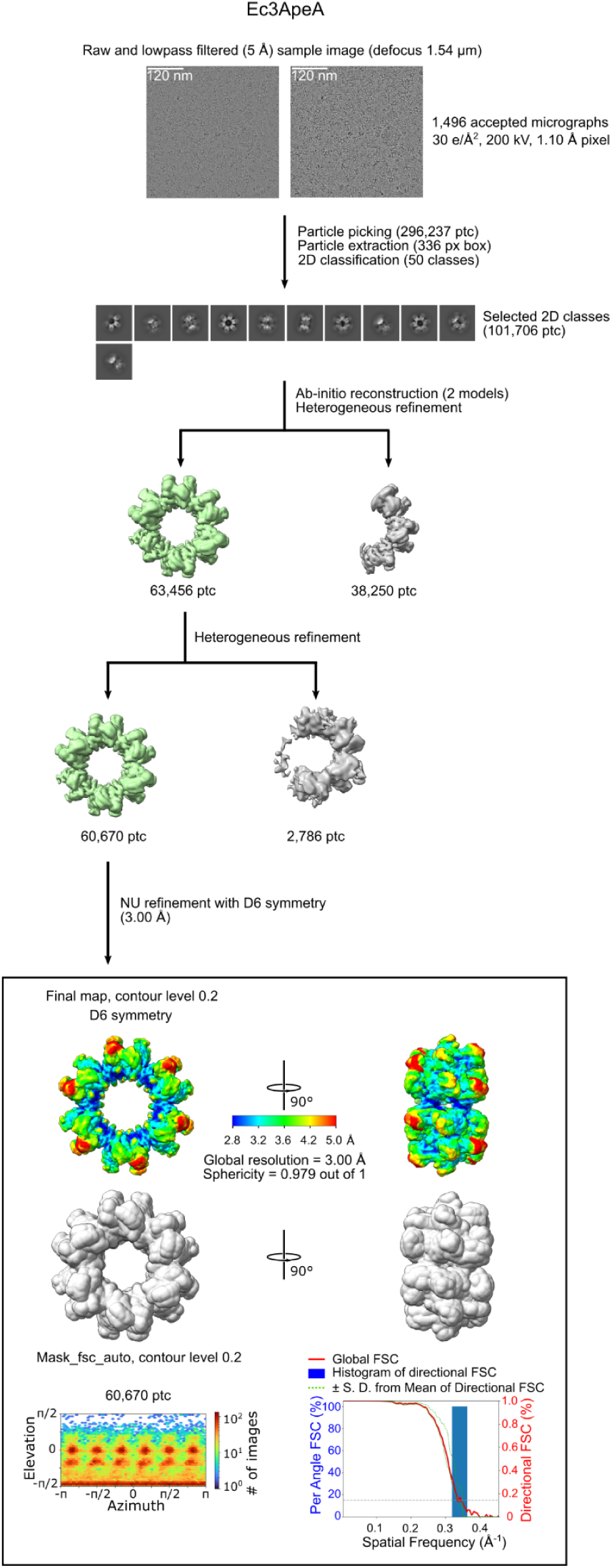
Cryo-EM data processing of Ec3ApeA.

**Figure S14.**
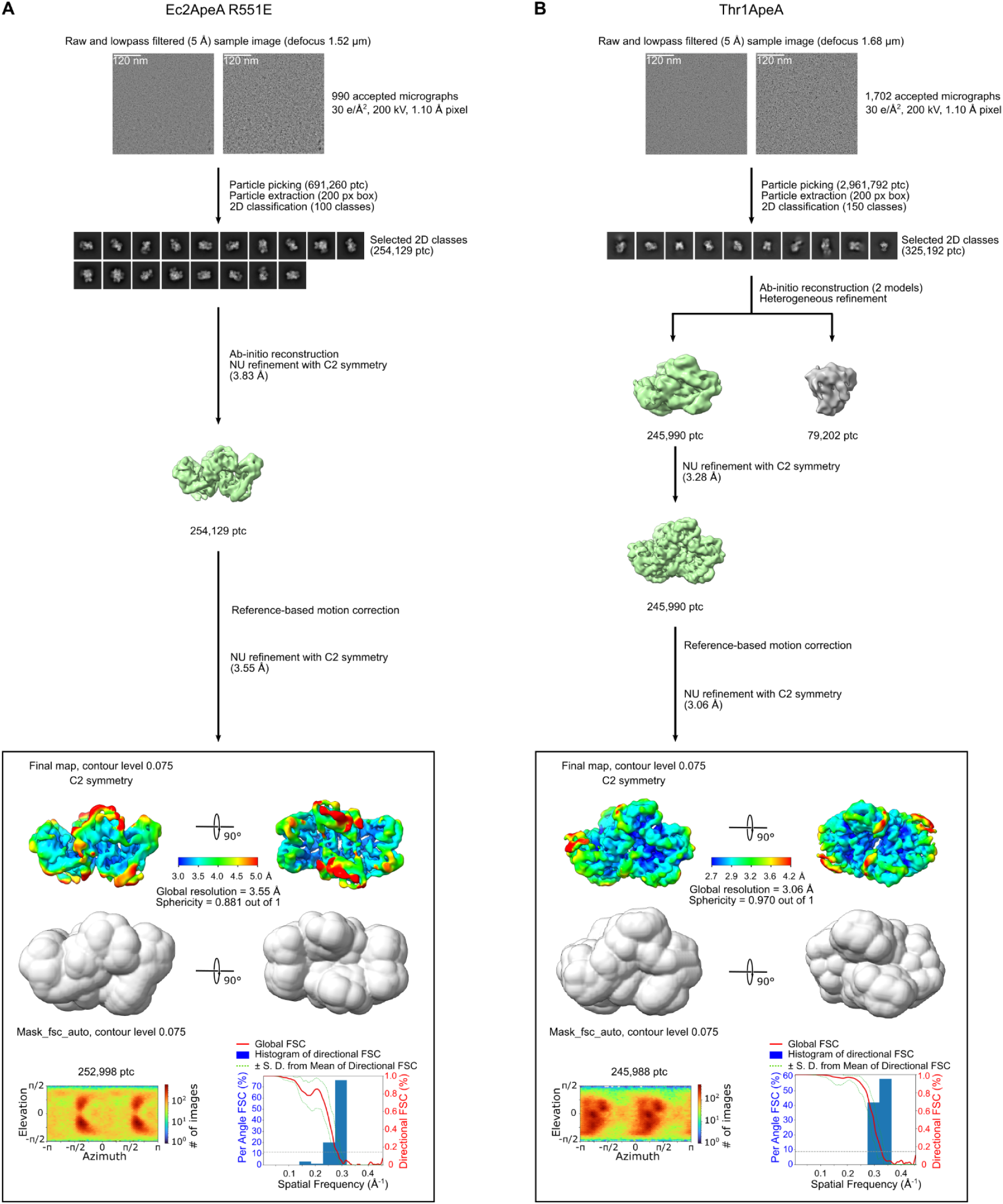
Cryo-EM data processing of Ec2ApeA R551E mutant (A) and Thr1ApeA (B).

**Supplementary Table S1.**
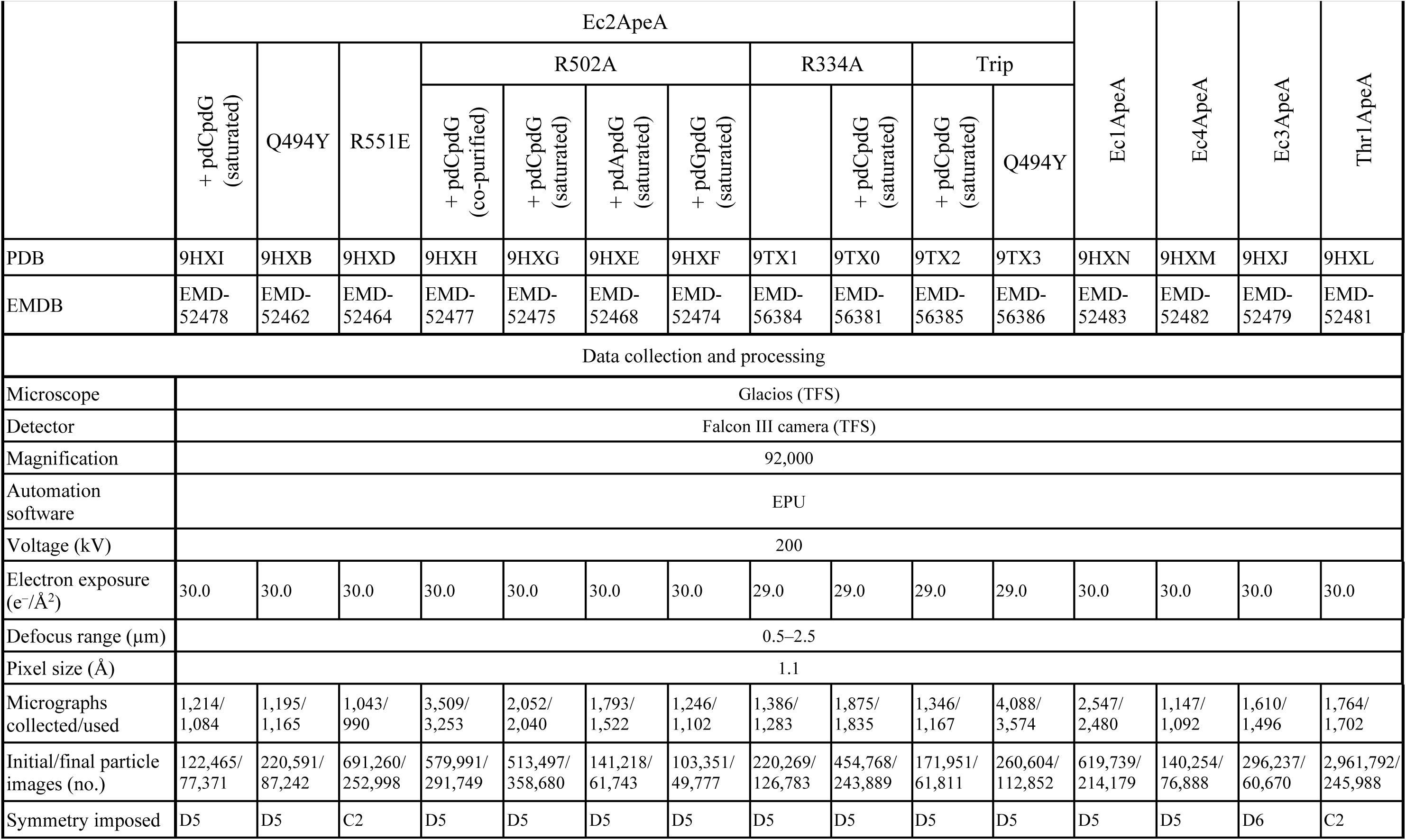
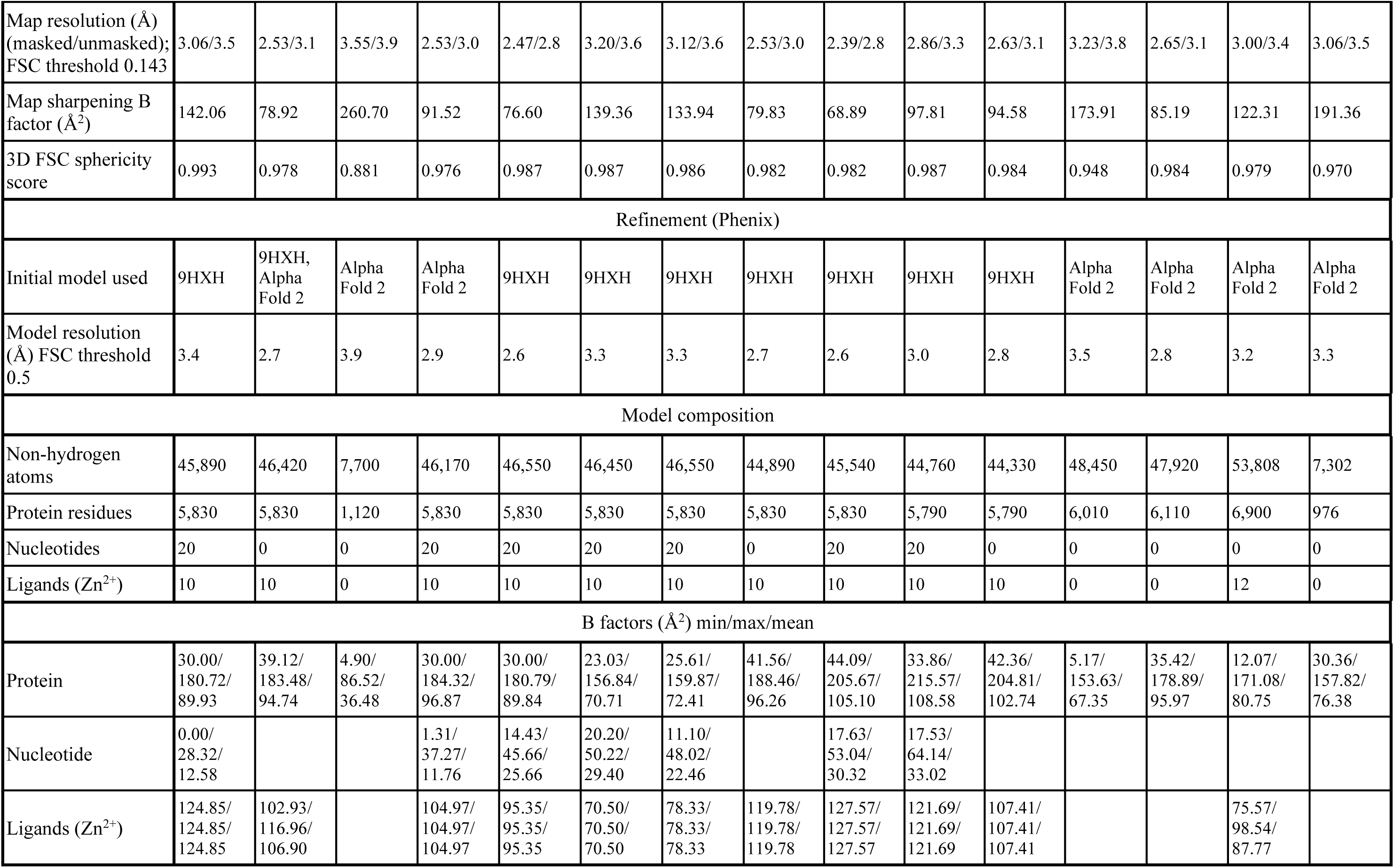
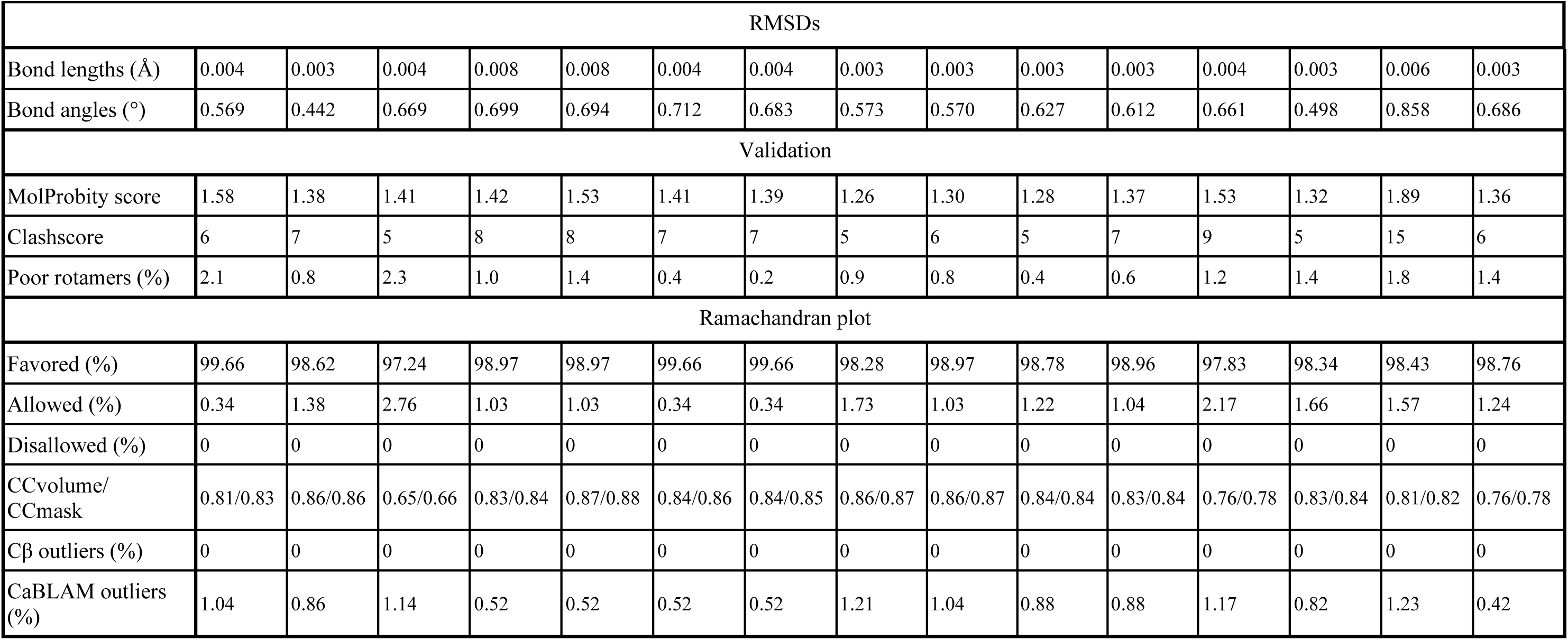
Cryo-EM data collection, refinement and validation statistics.

## METHODS

### Selection of ApeA proteins

Ec1ApeA (WP_000706972.1) was used as a starting query for 3 PSI-BLAST search itterrations using the nr database with the following parameters: Expect threshold: 1e-5, Word size: 3, Matrix: BLOSUM62, Gap Costs: existence 11, extension 1, PSI-blast Threshold: 0.01 ^43^. After each iteration sequences were filtered for query coverage >80% and identity <80%. After the third PSI-BLAST iteration sequences were clustered to 40% identity and 80% query coverage using MMseqs2 with clustering mode: normal, (version c552cce6c3194c06bc0bba84f04c4ef13d62f0a5). The resulting sequences were aligned using MAFFT (v7.273) algorithm (gap open penalty 1.53) on the MPI Bioinformatics tool kit server ^44^. The alignment was visualized using Ugene (v51.0) and manually curated by removing sequences without the HEPN active site residues, sequences with large insertions or deletions and proteins seemingly not fitting the alignment. Poorly aligned N and C sections of the alignment were removed, leaving the alignment spanning 21-557 residues of Ec1ApeA. The alignment was further clipped using clipkip (v2.3.0; --mode smart-gap) ^45^. Maximum likelihood phylogenetic tree was built using IQ-TREE2 ^46^ with automatic model selection and ultra-fast bootstrap approximation (v2.3.6; -B 10000). The phylogenetic tree was visualized using iTOL (v7.0) ^47^. ApeA variants for experimental analysis were selected from branches more distant from Ec1ApeA.

Sequences in the final set originating from RefSeq database records were further analysed by the WebFlags ^48^ web server (available at: https://server.atkinson-lab.com/webflags) to examine the genomic neighborhoods of ApeA proteins for functionally linked genes.

### Structure based sequence alignment

The set of ApeA protein sequences from the final manually curated sequence alignment were used to predict dimeric structures using the Colabfold implementation of Alphafold2 ^49^ generating 5 models and using the best one judged by plDDT metric. The similarity of predicted structures to the Ec2ApeA WT experimental cryo-EM structure was evaluated using GTalign v0.19.00 ^27^ and structures with TM-score <0.85 were discarded. Then structures of the single chain were extracted using a custom python script and the experimental monomeric structures of Ec1, Ec2, Ec3, Ec4 and Thr1 were added to the set, yielding 72 structures in total. Structure based alignment was performed using the foldmason v4.dd3c235 ^50^ “easy-msa” command.

### Cloning and mutagenesis

Genes were ordered as synthetic DNA fragments and cloned in a low-copy pACYC vector with an upstream IPTG inducible T7 polymerase promoter. The genome encoding Ec4 ApeA was later noted as possibly contaminated, from a large multi-isolate project, unverified source organism by the National Center for Biotechnology Information (NCBI). The Thr1ApeA gene was codon optimized for expression in *E. coli.* All ApeA variants except Ec3 ApeA and Pa1ApeA were fused with C-terminal His-tag for protein expression and purification. Additionally, Ec1ApeA and Ec3ApeA were cloned in a pTK120 vector with a TEV cleavable MBP-His10-twinStrep supertag using the Gibson assembly method. ApeA gene point mutant vectors were constructed by whole plasmid amplification using primers with 5′ mismatches introducing mutations followed by phosphorylation and blunt-end ligation. For the protection assay against the BASEL II phages ^21^ Ec2ApeA gene was cloned under the native promoter of Ec1, as the strain used for BASEL II phages BW25113 ΔgtrS wbbL(+) does not code T7 RNA polymerase. All constructs were sequenced to confirm the correct sequence.

### Phage propagation

All bacteriophages were propagated in *E. coli* MG1655 strain and BASEL II phages in BW25113 ΔgtrS wbbL(+). *E. coli* cells were grown in LB medium until OD_600_ reached 0.6-0.8, supplemented with 5 mM CaCl_2_ and 5 mM MgCl_2_, infected with phage at MOI 0.01-0.05, and incubated for 16 hours at 37 °C. Cultures were centrifuged for 10 minutes at 6000g and supernatants were filtered through 0.22 µm filters.

### Plaque assay

*E. coli* HMS-174(DE3) cells harboring the vector coding for the antiviral system from a streak plate were sampled and inoculated into growth medium, cultivated for 3-4 hours, and induced with 0.05 mM IPTG if needed (depending on the experiment). BW25113 ΔgtrS wbbL(+) strain for the infection with BASEL II phages was not induced. The solid layer of LB medium with 1.5% agar was supplemented with the appropriate antibiotic. The soft layer of LB medium with 0.5% agar (soft agar layer) was supplemented with the appropriate antibiotic, inducer (for HMS174(DE3); 0.05 mM IPTG or 0.05% arabinose), 5 mM CaCl₂, and 5 mM MgCl₂. Two hours after the induction, 200 µL of cell culture were added to the soft agar, gently mixed, and immediately poured onto the first layers of agar. The plates were kept open to dry for 15 minutes. Then, 3.5 µl of the serially diluted phage stock suspension were spotted on the bacterial lawns, open plates were kept open for 10 minutes for the spotted solution to dry and absorb. Plates were incubated for 16 hours at 37°C. The lowest dilution with easily counted plaques was used to estimate the phage titer. The ratio between the number of plaques formed on the control and ApeA-containing lawn (accounting for the dilution factor) was calculated and referred to as the *protection ratio*.

Ec1, Ec2, Ec4, Thr1 had a fused C-terminal His-tag, while Ec3 and Pa1 did not have the tag in plaque assay experiments. Presence of C-terminal His-tag did not interfere with protein activity *in vivo*. Ec1 and Ec2 ApeA homologs were not tagged in the plaque assays of BASEL II phages.

### Protection assay in liquid medium

*E. coli* HMS-174(DE3) were transformed with vectors coding His-tagged Ec1 and Ec2 ApeA proteins and a control vector without the insert. 200 µL night cultures were transferred to 4 ml LB. Cells were grown until OD_600_ reached 0.6, induced with 0.05 mM IPTG and grown for 3 hours, at 37 °C, 200 rpm shaking. The culture was diluted to OD_600_ 0.3 and infected with phage RB69 or Bas27 at a MOI of 0.05 and 10 in the final volume of 100 µl in a 96-well plate. The inside of the plate lid was treated with 20% ethanol 0.05 % Triton-X100 solution and dried to stop the formation of water droplets^51^. The sides of the plate were sealed with parafilm and the plate was incubated at 37 °C with shaking at 200 rpm. Optical density was measured every 5 minutes.

### Toxicity assay in liquid medium

*E. coli* HMS-174(DE3) cells were transformed with a pBAD vector carrying the diDNase gene variants and pACYC vector coding either Ec1 or Ec2 His-tagged ApeA protein variants. 200 µL of overnight culture was inoculated into 4 mL of LB. Cultures were grown in LB medium supplemented with the appropriate antibiotics at 37 °C with shaking at 200 rpm until an OD_600_ of 0.6 was reached. Expression of diDNase was induced with 0.05% arabinose, and cultures were incubated for an additional 3 h.

Cells were next diluted to an OD_600_ of 0.1 in LB medium containing appropriate antibiotics, 0.05% arabinose and 0.1 mM IPTG to induce the expression of ApeA proteins. Diluted cells were dispensed into a 96-well plate at a final volume of 100 µL per well. The plate lid was treated with 0.05% Triton X-100 and 20% ethanol to prevent condensation. The closed plate was sealed with parafilm and incubated at 37 °C with shaking at 200 rpm. Optical density was recorded every 5 minutes.

### Protein expression and purification

#### Protein expression

*E. coli* BL21 (DE3) was used for the expression of ApeA proteins cloned in pACYC under IPTG inducible T7 promoter, while the BL21-AI strain was used for the expression of proteins in pTK120 vector under arabinose inducible araBAD promoter. Bacterial cultures were grown at 37 °C. When the OD_600_ reached 0.5-0.8, the culture was cooled to 4-10 °C; the cold culture was induced with 0.25 mM IPTG (T7 promoter) or 0.05% arabinose (araBAD promoter) and incubated overnight at 16 °C. Cells were collected by centrifugation and the pellet was stored at -20 °C.

#### Purification of His-tagged proteins

Cells were thawed, resuspended in buffer A (20mM Tris-HCl, pH 8, 500 mM NaCl, 5 mM 2-mercaptoethanol), supplemented with Halt Protease Inhibitor Cocktail (Thermo Scientific), and disrupted by sonication, with the sonication container submerged in ice bath. The lysate was centrifuged at 18000 g for 1 hour at 4 °C, supplemented with 2% (v/v) buffer BH (20mM Tris-HCl, pH 8, 500 mM NaCl, 5 mM 2-mercaptoethanol, 500 mM imidazole) and the supernatant was subjected to Ni^2+^ affinity chromatography purification through 5 ml HisTrap HP (Cytiva) column. After sample loading, the column was washed with 10 column volumes (CV) of A buffer. The protein was eluted with 10 CV of 2-100% buffer BH gradient. Protein fractions containing the protein of interest were pooled, concentrated by centrifugation (10K column concentrator), and subjected to SEC on a HiLoad 16/600 Superdex 200 (Cytiva) column. Fractions containing the protein of interest were concentrated by centrifugation and dialyzed against a storage buffer S (20 mM Tris-HCl, pH 8, 500 mM NaCl, 2 mM DTT, 50% (v/v) glycerol).

#### Purification of MBP-His-Twin-Strep-tagged proteins

The bacteria pellet for MBP-Ec1ApeA purification was resuspended in the same buffer A, but with a higher salt concentration - 1.5 M, to promote a more efficient dissociation of unspecifically bound nucleic acids. Cells were lysed and processed following the same protocol as for His-tagged proteins and subjected to 5 ml StrepTrap HP (Cytiva) column. The column was washed with 10 CV of buffer A and the protein eluted with 5 ml of buffer BS (20mM Tris-HCl, pH 8, 500 mM NaCl, 5 mM 2-mercaptoethanol, 2.5 mM desthiobiotin). Fractions containing the protein of interest were pooled and concentrated by dialysis against buffer A with 50% glycerol for the second purification step by SEC. All other purification procedures were the same as for His-tagged proteins. The desired fractions were pooled and dialyzed against the buffer S.

### Mass spectrometry of protein bound ligands

#### Sample preparation

The purified proteins were diluted to 2 mg/ml with water and denatured by heating at 80 °C for 10 minutes. Samples were centrifuged at 12000 g for 30 min to separate protein aggregates and the solution was subjected to mass spectrometry.

#### HPLC/ESI-MS

Samples were analyzed on an integrated HPLC/ESI-MS system (Agilent 1290 Infinity/ Agilent Q-TOF 6520 mass analyzer) equipped with a Supelco Discovery® HS C18 column (7.5 cm × 2.1 mm, 3 µm). Mass analyzer was operating in the positive ion mode. Elution was performed with a linear gradient of solvents A (0.02% formic acid in water) and B (0.02% formic acid in acetonitrile) at a flow of 0.3 ml/min at 30 °C as follows: 0-5 min, 0%B; 5-18 min, 0-14%B; 18-22 min, 14-95 %B; 22-27min, 95%B.The results were analyzed with Agilent MassHunter Qualitative Analysis software B.05.00.

#### HPLC/ESI-MS/MS

Samples were analyzed on an integrated HPLC/ESI-MS/MS system (Agilent 1290 Infinity/ 6410B triple quadruple) equipped with a Supelco Discovery® HS C18 column (7.5 cm × 2.1 mm, 3 µm). HPLC conditions were the same as in the case of HPLC/ESI-MS (as described above). Mass analyzer was operating in the positive ion MRM mode and intensity of nucleotide-specific ion transition was recorded: 637.1→152.1 (dpCG→G) and 637.1→112.1 (dpCG→C).

### Detection of cellular dinucleotides

#### Sample preparation

*E. coli* HMS-174(DE3) cells were transformed with a pBAD expression vector carrying the diDNase gene or with an empty pBAD vector as a control. 1 mL of overnight culture was inoculated into 50 mL of LB medium supplemented with the appropriate antibiotic and grown at 37 °C with shaking at 200 rpm until an OD_600_ of 0.6 was reached. Protein expression was induced with 0.05% arabinose for 1 h, after which OD_600_ was measured again. Cells were collected by centrifugation at 6000 g for 2 min, the supernatant was discarded and the pellets were immediately frozen in liquid nitrogen.

Cell pellets were resuspended in methanol at a volume proportional to culture density (400 µL per a single unit of OD_600_). Cell lysis was achieved by three freeze–thaw cycles (freezing in liquid nitrogen and thawing on ice). Lysates were centrifuged at 12000 g for 60 min at 4 °C. The resulting supernatant was heated at 75 °C for 10 min and centrifuged again at 12000 g for 30 min at 4 °C. Supernatants were transferred to low-binding microcentrifuge tubes and concentrated using a vacuum concentrator at 45° C. Concentrated samples were diluted with MiliQ water in proportion to the initial culture density (100 µL of water per a single unit of OD_600_).

### Preparation of phosphorylated dinucleotides

Dinucleotides were synthesized as desalted oligonucleotides and phosphorylated enzymatically using T4 polynucleotide kinase (Thermo Scientific). 500 µM of dinucleotide and 1 mM ATP were diluted in the PNK buffer (50 mM Tris-HCl pH 7.6, 10 mM MgCl_2_, 5 mM DTT, 0.1 mM spermidine). 30 µl reactions were supplemented with 1 µl of 10 U/µl PNK and incubated for 1 hour at 37 °C. After incubation, the reactions were inactivated by heating at 80 °C for 10 minutes and centrifuged for 30 minutes at 12000 g to remove the denatured proteins. The supernatant was stored at -80 °C. An identical reaction without dinucleotides was prepared as an ATP control.

### Oligomeric state determination by SEC-MALS

Size-exclusion chromatography with multi-angle light scattering (SEC-MALS) of all proteins complexes (200 μl sample, 5 μM of protein monomer) were carried out at room temperature using a Superdex 200 Increase 10/300 GL column (Cytiva) equilibrated with buffer (20 mM Tris–HCl (pH 8.0 at 25 °C), 150 mM NaCl, 2 mM MgCl2, 1 mM DTT), at 0.4 ml/min flow rate. The light scattering signals were monitored on a miniDawn TREOS II detector, concentrations of all samples were measured using an Optilab T-rEX refractive index detector (Wyatt Technologies). Data were analysed in Astra software (Wyatt Technologies) using a dn/dc value of 0.185 ml/g.

### Sample preparation for Cryo-EM

The protein fraction with the highest concentration, identified at the center of the size exclusion chromatography (SEC) elution peak, was concentrated using Amicon Ultra 0.5 mL 10K concentration columns (Merck Millipore). When needed, the proteins were diluted in CE buffer (20 mM Tris-HCl, pH 8, 500 mM NaCl, and 1 mM DTT) to achieve a final concentration of 2 mg/mL (∼30µM monomer). For samples requiring a dinucleotide, 100 µM of the phosphorylated dinucleotide (∼3x the protein monomer concentration) was added. The mixture was incubated at room temperature for 10 min before being transferred to 0 °C. 3 µl of the sample was applied to a glow-discharged (20 mA for 45 s) Quantifoil Cu grids (300 mesh, R 1.2/1.3) and vitrified in liquefied ethane using Vitrobot Mark IV (FEI) at 4 °C and 100% humidity with 0 s waiting and 6 s blotting times.

### Cryo-EM data collection and image processing

Cryo-EM data collection was performed using a Glacios microscope (Thermo Fisher Scientific), operating at 200 kV and equipped with a Falcon 3EC Direct Electron Detector in the electron counting mode (Vilnius University, Vilnius, Lithuania). Data was collected with EPU (v.2.11.0 - v.3.13) at a nominal magnification of 92000×, corresponding to a calibrated pixel size of 1.10 Å, using an exposure of 0.80 e^−^/Å^2^ s^−1^, in 30 frames and a final dose of 29-30 e^−^/Å^2^, over a defocus range of −1.0 to −2.0 µm. Cryo-EM data collection is summarized in Supplementary Table S1.

Patch motion correction, CTF estimation, micrograph curation and, in some cases, blob picking with particle extraction were performed in real-time in CryoSPARC Live v.3.2.0 - v.4.7.1 ^52^. Further data processing and final refinement were performed using standard CryoSPARC v.3.2.0 - v.4.7.1 ^52,53^. Image processing for each sample is summarized in (Supplementary Figures S7-14). In all cases, 3DFSC web server ^54^ was used to estimate the global resolution at 0.143 FSC cutoff and sphericity values of the final electron density maps, while the local resolution was calculated using CryoSPARC (v.4.7.1).

### Cryo-EM model building, refinement and analysis

Initial models of Apea dimers were generated using AlphaFold 3 ^55^. Using ChimeraX (v.1.9) ^56^, an appropriate number of dimeric structures were fitted into the Cryo-EM map based on the oligomeric assembly of each ApeA protein. Protein rebuilding and manual building of dinucleotides were performed using Coot (v.0.9.8.1) ^57^. Models were refined using phenix.real_space_refine (v. 1.21.2-5419) ^58^, refinement statistics are summarized in Supplementary Table S1. Structure overlays and generation of structural images were performed using ChimeraX (v.1.9).

### Differential Scanning Fluorimetry

Proteins were diluted to 7.25 µM in DSF buffer (33 mM Tris-HCl pH 7,9, 66 mM K-acetate, 200 mM NaCl, 10 mM MgCl_2_, 0.1 mM DTT) and mixed with 50 µM of the phosphorylated dinucleotide or ATP control. Mixtures were incubated at room temperature for 10 minutes prior to the experiments. Protein melting temperatures were assessed on the Nanotemper Prometheus device, using 1 °C/min temperature increments. For every reaction, 2 replicates were prepared. Data was analyzed using the Prometheus NT.48 software.

### RNA extraction

200 µl of *E. coli* BL21 (DE3) overnight culture with a plasmid expressing WT Ec1ApeA system or the deactivated Ec1ApeA control were used to inoculate 4 ml of LB medium. The cultures were grown for 1 hour at 37 °C, induced with 0.1 mM of IPTG and incubated for 3 more hours until OD_600_ reached 0.5-0.8k, and then were infected with bacteriophage T2 at a MOI of 5. 1 ml of bacterial cultures at different time points (0, 10, 20, 45, 60, 90 minutes) after infection were centrifuged for 1 min at 12000 g; the supernatant was removed and the cells were immediately frozen in liquid nitrogen. After collection of all the samples, the cells were resuspended in 100 µl of TE buffer (10 mM Tris HCl, pH 8.0, 1 mM EDTA) supplemented with 0.4 mg/ml lysozyme and incubated for 2 minutes at 37 °C. RNA was extracted using Thermo Scientific GeneJET RNA Purification Kit following the Bacteria Total RNA Purification protocol. 300 ng of RNA were run on 10% 29:1 Urea-PAGE and stained with SYBR Gold.

### RNA sequencing and data analysis

#### Library preparation

1 µg of isolated RNA was incubated with 1 µl of 1 U/µl DNase I (Thermo Scientific) in 50 µl DNase I reaction buffer (10 mM Tris-HCl pH 7.5, 2.5 mM MgCl_2_, 0.1 mM CaCl_2_) for 30 minutes at 37 °C. 10 mM ATP and 2 µl of 10 U/µl PNK were added to the reaction, and the volume diluted to 100 µl using 1x PNK A buffer (50 mM Tris-HCl pH 7.6, 10 mM MgCl_2_, 5 mM DTT, 0.1 mM spermidine). The reaction was incubated at 37 °C for another 30 minutes. After incubation, 10 mM EDTA was added and the enzymes were deactivated by incubation at 75 °C for 10 minutes. The RNA was split into small and large RNA fractions using the “Lexogen Split RNA extraction kit” following the “Phenol/Chloroform Extraction” and “Column-based Purification” protocols. The large RNA fractions were used for sequencing. Sequencing libraries were barcoded and prepared using the “Lexogen Small RNA-Seq Library Prep Kit” according to the manufacturer’s instructions and sequenced on the Illumina Miniseq platform using 2×150 paired-end mode.

#### Data analysis

Adapters were trimmed and filtered using BBduk (ktrim=r k=23 mink=10 hdist=1 tpe tbo minlen=17), from the BBTools package (version 38.18, https://sourceforge.net/projects/bbmap/) aligned to BL21 (DE3) reference genome using bowtie2 under default parameters, filtered (-F 4) to include only unambiguously mapped reads and sorted by samtools (v1.12)^59^. Read numbers were normalized according to the dataset with the least reads. Data was analyzed using custom Python scripts. In brief, the number of read ends mapping to the position was counted at each genomic coordinate. The positions where the read end count ratio of normalized “active ApeA”/”inactive ApeA” datasets was the highest, were selected for further inspection. A cutoff threshold of 100 was used. Genomic features mapping to the positions with selected ratios were analyzed and visualized using custom Python scripts.

### Structure analysis

*General analysis* and structure visualization was performed using ChimeraX v1.9^56^. Structures homologous to ApeA domains were found using DALI webserver ^60^ hosted at http://ekhidna2.biocenter.helsinki.fi/dali/ and GTalign webserver, hosted at https://bioinformatics.lt/comer/gtalign/^27,41^, with default parameters. The structures with the most aligned residues were selected for further analysis and curation.

*Conservation analysis* was conducted using the Consurf ^42^ webserver (available at: https://consurf.tau.ac.il/). Residues for ligand binding pocket mutations were selected by manually analyzing the structures and based on the Consurf conservation score.

*Oligomerization interface analysis.* Oligomerization interfaces were analyzed using the PDBePISA server (Protein interfaces, surfaces and assemblies’ service PISA at the European Bioinformatics Institute; http://www.ebi.ac.uk/pdbe/prot_int/pistart.html) with default parameters. Interface residues for mutagenesis were selected based on structural examination.

Conformational changes were analysed using UCSF ChimeraX (v1.9) by global alignment of two structures using the “matchmaker” function by associating the structures by sequence, and rendering the “RMSD C-ɑ” attribute for each residue. The changes in contacts were also analysed using the VoroContacts (available at: https://bioinformatics.lt/wtsam/vorocontacts) online webserver ^61^ by manually comparing the contacts of each structure.

### tRNA cleavage assay

ApeA proteins were assayed in the buffer R: 33 mM Tris-HCl, pH 7.9, 66 mM KOAc, 1 mM DTT, 0.5 U/µl RiboLock RNAse inhibitor (Thermo Fisher Scientific). tRNA concentration was 1 µM, and protein concentration (monomer) 1 µM, ligand concentration was 50 µM. The final reaction volume was 10 µl. The reaction was started by adding 1 µl of 10x concentrated protein in buffer R to the mixture of other components. Reactions were incubated at 37 °C for 90 min and stopped by adding 10 µl of 2x RNA loading dye (Thermo Fisher Scientific). Samples were heated for 5 minutes at 80 °C and immediately transferred on ice. Samples were run on 15% Urea PAGE in TBE buffer (acrylamide:bis-acrylamide ratio 1:29), 5 V/cm, and stained with SYBR Gold.

