## Supplementary File S2.pdf for "Deoxydinucleotides activate the bacterial anti-phage defense system ApeA"

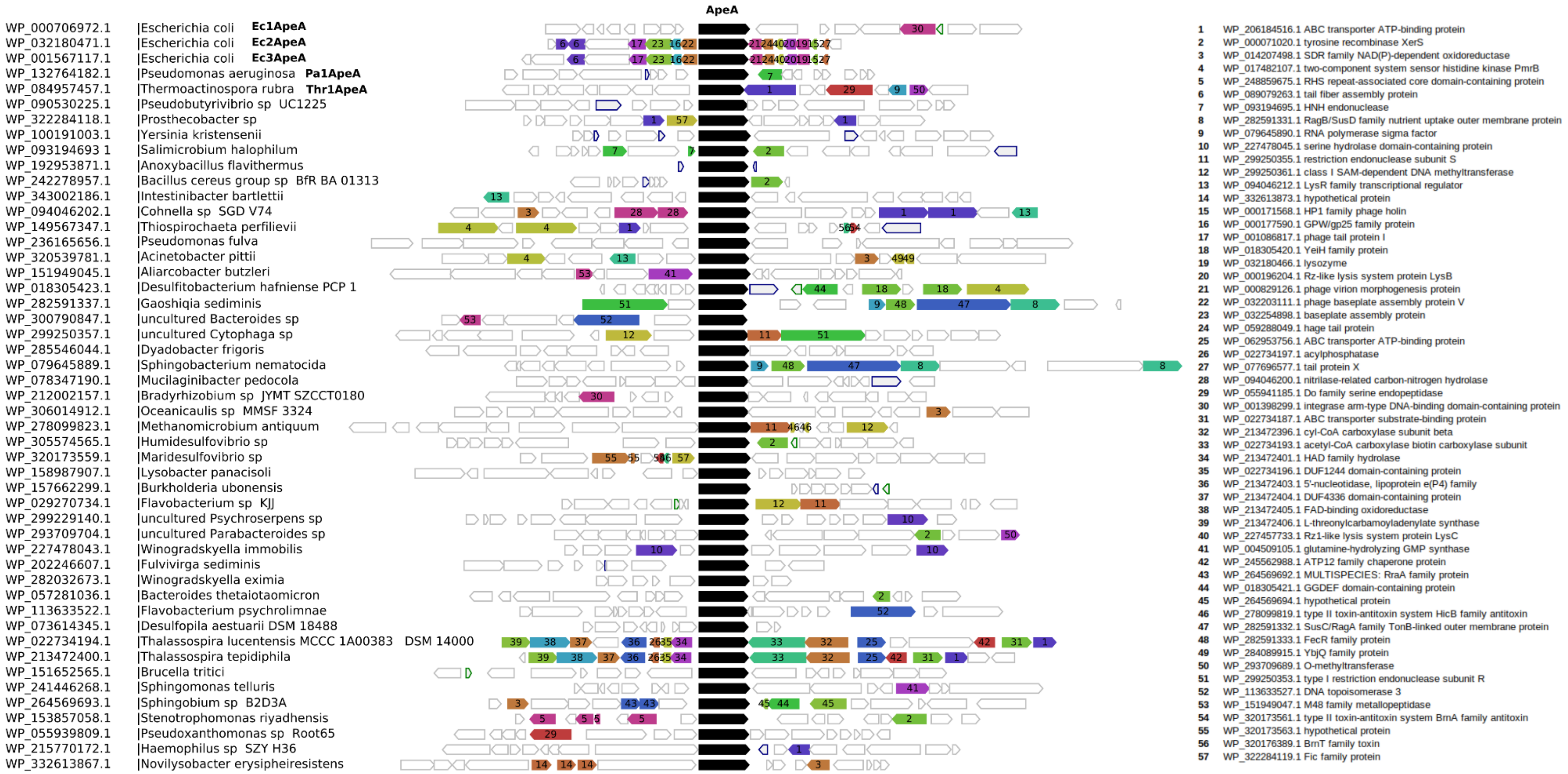


WebFlags analysis of ApeA sequences from RefSeq database (2025-12-12) used to build ApeA phylogenetic tree of Figure 1. Proteins selected for functional characterization marked in bold. Only sequences from RefSeq database were used for the analysis. ApeA is  a standalone defense system, not functionally linked to other genes. Only ApeA protein genres were selected for cloning.
